# The Deacetylases HDAC1 and HDAC2 Safeguard BCR-ABL-positive Cells from Replication Stress-Induced Apoptosis via the Nuclear to Mitochondrial p73-NOXA Axis

**DOI:** 10.64898/2025.12.13.694110

**Authors:** Al-Hassan M. Mustafa, Miriam Pons, Nisintha Mahendrarajah, Maren Wiegerling, Jörg Hartkamp, Markus Christmann, Markus P. Radsak, Günter Schneider, Matthias Wirth, Oliver H. Krämer

## Abstract

Histone deacetylases (HDACs) are key epigenetic regulators that are frequently dysregulated in cancer cells. Context-specific dependencies of tumor cell survival on HDACs and the mechanisms through which HDACs determine cell stress responses by specific oncogenic pathways remain to be understood. Here we unravel how the class I deacetylases HDAC1 and HDAC2 control the fate of chronic myeloid leukemia (CML) cells undergoing DNA replication stress. Compared to normal myeloid cells (n=690), CML cells overexpress HDAC1 and HDAC2 (n=234-274). We reveal that HDAC1 and HDAC2 protect cultured and primary CML cells with the hyperactive BCR-ABL tyrosine kinase from programmed cell death through apoptosis upon DNA replication stress. Using transcriptomics and genetically defined knockdown and knockout model systems, we demonstrate that these effects depend on the transcription factor p73 and its pro-apoptotic target gene *NOXA*. Upon DNA replication stress, p73 binds to the *NOXA* promoter but only upon additional inactivation of HDAC1/HDAC2 the *NOXA* gene becomes transcribed. BCR-ABL translocate to the nucleus to catalyze p73 phosphorylation at tyrosine-99 for the induction of p73 and mitochondrial NOXA. Thus, the BCR-ABL oncogene creates a selective vulnerability to HDAC1/HDAC2 inhibitors, driving a cytotoxic shift in replication stress responses towards apoptosis. These data highlight HDAC1/HDAC2 as potential therapeutic targets in CML.

**Graphical Abstract:** 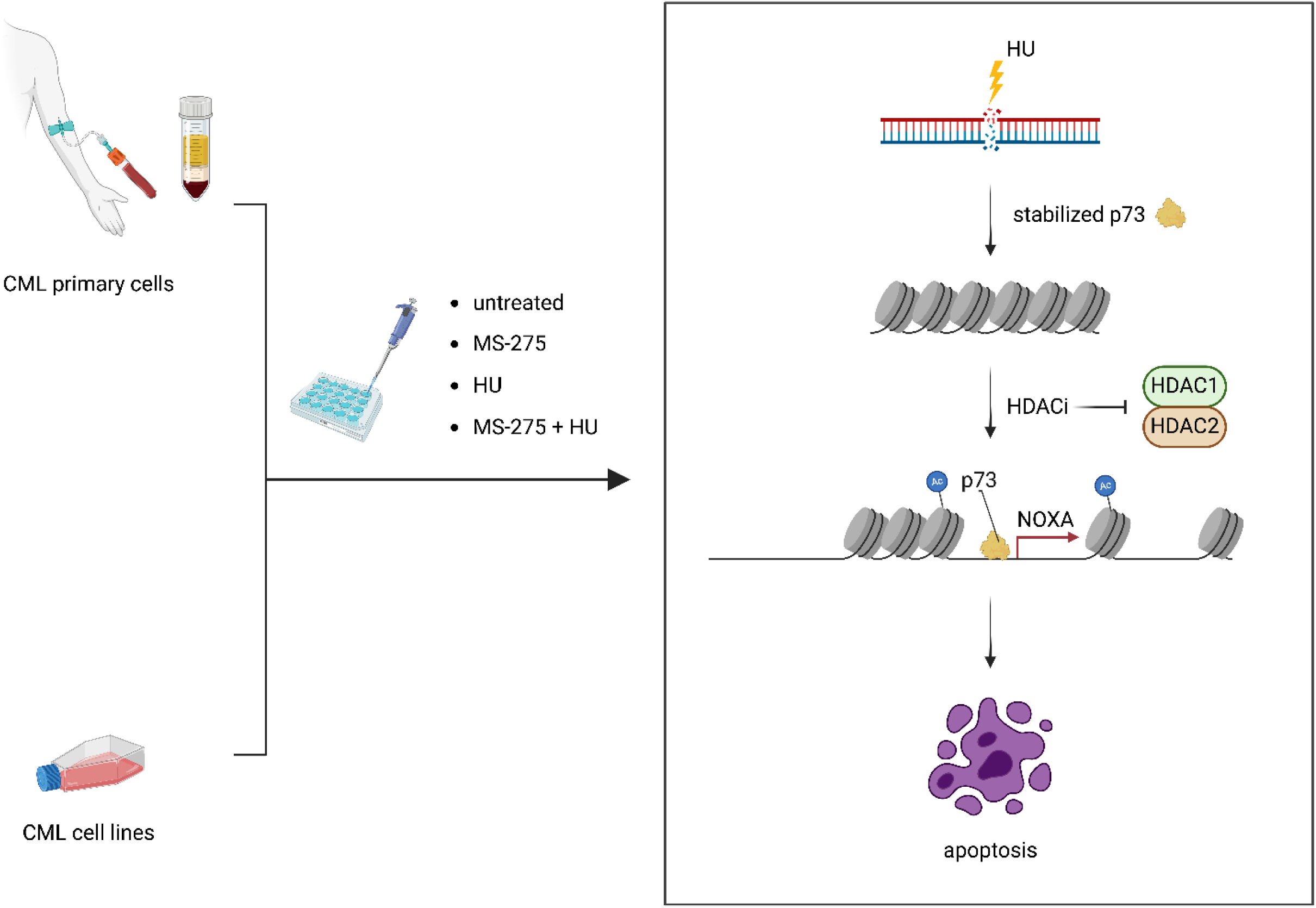

## Introduction

Apoptosis, a form of regulated cell death, is a tightly controlled process that eliminates unnecessary, damaged, or transformed cells. This makes apoptosis one of the most critical barriers to tumorigenesis (Sarosiek and Wood, 2023). Apoptosis is orchestrated through the intrinsic (mitochondrial) and extrinsic (death receptor) pathways. Both culminate in the activation of caspases that dismantle cellular proteins and lead to chromatin fragmentation by caspase-activated DNase. The release of cell fragments by apoptotic cells allows their uptake by neighboring cells and an avoidance of inflammation that occur upon uncontrolled cell rupture (necrosis). Central to the intrinsic apoptosis pathway, which is triggered by most if not all frequently applied chemotherapeutics, are B cell lymphoma-2 (BCL2) proteins. These include pro-apoptotic and anti-apoptotic members (Shamas-Din et al., 2011). NOXA belongs to the pro-apoptotic BH3-only proteins that localize predominantly to mitochondria. NOXA promotes permeabilization of the mitochondrial outer membrane by the pore-forming BCL2 family members BAX and BAK. The following efflux of cytochrome c triggers caspase-9 activation to initiate apoptosis via the ultimate apoptosis executioner caspase-3 (Shamas-Din et al., 2011). Despite such known functions of NOXA, its context-specific dependencies are incompletely charted. If NOXA acts beyond a passive apoptotic trigger and creates vulnerabilities when combined with specific oncogenic pathways (e.g., translocation of the leukemia fusion protein BCR-ABL, MYC overexpression, RAS mutations) likewise remains to be revealed.

Optimal chemotherapy exploits key features of cancer cells. These include a high proliferation index and a resulting excessive demand for metabolites and nucleotides as well as a typical signature of mutations (Sarosiek and Wood, 2023; Shang et al., 2024). The unrestricted proliferation rates of tumor cells create a window for pharmacological strategies targeting DNA replication. The term DNA replication stress describes an obstruction of the processivity of DNA replication forks. This can occur for example, due to excessive DNA replication and dNTP shortage, and upon an insertion of nucleoside analogs in DNA that inhibit DNA polymerase processivity. Hydroxyurea (HU) is frequently used to cause DNA replication stress in cultured cells and is a first-line treatment for blood cancers (Dzulko et al., 2020). HU inhibits the ribonucleotide reductase M2 (RRM2) subunit of the ribonucleotide reductase complex and consequently impairs de novo dNTP synthesis. This slows down DNA polymerases and causes an accumulation of single strand DNA (ssDNA) stretches due to an ongoing activity of replicative minichromosome maintenance DNA helicases. Replication protein A (RPA) covers and protects ssDNA. Upon prolonged DNA replication stress, replication forks are destabilized, collapse, and eventually form ssDNA breaks and DNA double strand breaks (DSBs) (Göder et al., 2018; Nguyen et al., 2021). Cytarabine (1-β-D-arabinofuranosylcytosine) is also a major chemotherapeutic agent. This analog of deoxycytidine contains arabinose instead of deoxyribose and becomes phosphorylated inside the cell to its active form cytarabine triphosphate. During S-phase, cells incorporate cytarabine triphosphate into DNA, such sites stall DNA polymerases, cause DNA chain termination, and collapsing DNA replication farks that become DNA breaks (Gielecinska et al., 2023; Park et al., 2020). The experience with HU and cytarabine for decades encourages investigations to fully exploit and optimize their anti-cancer effects.

Chemotherapy may be improved through the identification of molecular mechanisms that shift tumor cells with DNA replication stress into apoptosis. Such mechanisms include the activation of the tumor-suppressive transcription factors p53 and p73. These induce apoptosis and other types of cell death (Cai et al., 2022; Rozenberg et al., 2021). Structural similarities of p53 and p73 are found in their DNA binding domains. This translates into a significant overlap in their target genes. Among them are genes that encode proteins promoting cell cycle arrest and apoptosis, such as NOXA. Since p53 is often lost in tumor cells, remaining functions of p73 may compensate p53-directed gene subnetworks (Cai et al., 2022; Rozenberg et al., 2021). Whether p73 can fully replace p53, the context-specific nature of such functional overlap, and how p73 can be pharmacologically exploited in cells undergoing genotoxic stress has not been fully elucidated. Likewise, tumor-specific regulators of NOXA expression beyond p53, p73, and other transcription factors are mapped incompletely. This likewise applies for the functional relevance of the p73-NOXA signaling axis and if this molecular pathway affects cancer subtypes differentially.

Within this century, accumulating evidence has indicated that epigenetic modifiers of the HDAC family determine the cellular outcome of stress responses and that these enzymes are frequently dysregulated in pre-cancerous stages and tumors (Mustafa and Krämer, 2023). This corresponds to their impact on genomic and non-genomic regulatory levels. HDACs fall into four classes (I, IIa/IIb, III, IV) that catalyze the removal of acetyl groups from lysine residues in histones and non-histone proteins (Narita et al., 2019). This posttranslational modification consequently affects many processes, including cell cycle control and apoptosis, through modulatory effects on transcription, cellular signaling, and protein stability. At least 15 HDAC inhibitors (HDACi) are tested in clinical trials and Food-and-Drug-Administrations approved the HDACi vorinostat (SAHA), romidepsin (FK228), belinostat (PXD101), and chidamide for the treatment of subtypes of lymphoma and multiple myeloma. These HDACi are active against the 11 zinc-dependent HDACs (classes I, II, IV). According to current knowledge, HDACi with increased specificity may solve the problem of undesired effects that pan-HDACi cause (Mustafa and Krämer, 2023; Mustafa et al., 2025). Through an inhibition of anti-apoptotic proteins, an induction of pro-apoptotic proteins, and a modulation of DNA repair proteins, HDACi increase the efficacy of drugs that induce DNA replication stress and DNA damage. The class I HDACs HDAC1, HDAC2, and HDAC3 contribute to the maintenance of chromatin structure and genomic stability (Nikolova et al., 2017). Coherent with this notion, a combined application of structurally divergent HDACi (VPA, MS-275, LBH589) with HU evoked apoptosis of solid tumor cells in vitro and in vivo (Mustafa and Krämer, 2023; Nikolova et al., 2017). Cultured leukemia cells and leukemia cells in patients are also sensitive to combinatorial treatment with HU plus HDACi (Bug et al., 2005; Leitch et al., 2016), however the underlying molecular mechanisms are ill-defined. This notion and the approval of HDACi for the treatment of leukemia subtypes by Food-and-Drug-Administrations suggest further analyses on how individual HDACs control leukemia cell fate.

This work maps the transcriptional regulation of NOXA by p73 and its biological relevance upon DNA replication stress. We reveal that HDAC1 and HDAC2 suppress the p73-NOXA signaling node to protect chronic myeloid leukemia (CML) cells from apoptosis upon DNA replication stress. The oncoprotein of such cells is the BCR-ABL leukemia fusion protein arising from the chromosomal translocation t(9;22). Intriguingly, we show that BCR-ABL activity is a requirement of apoptosis induction by the p73-NOXA signaling node upon DNA replication stress and HDAC inhibition. This illustrates the first pro-apoptotic role of endogenous BCR-ABL and how it is targetable by targeted epigenetic drugs in cells with DNA replication stress.

## Material and Methods

### Permanent cell cultures and culture conditions

The BCR-ABL (translocation t(9;22)) positive e14-a2 CML cell line K562 was isolated from the pleural effusion of a 53-year-old female patient, and the erythroleukemia cell line HEL was from the bone marrow of a 30-year-old, white, male patient (both are kind gifts from Dr. Manuel Grez, Georg Speyer Haus, Frankfurt/Main, Germany). The CML cell lines LAMA-84 cells were collected from a 29-year-old female in CML blast crisis (4 copies of t(9;22)(q34;q11), e14a2 BCR-ABL), MEG-01 cells were from the bone marrow cells of a 55-year-old male in megakaryoblast CML (t(9;22)(q34;q11), e13a2 BCR-ABL), KCL-22 came from the pleural effusion of a 32-year-old female with CML in blast crisis (t(9;22)(q34;q11), e13a2 BCR-ABL), and KYO-01 cells were obtained from the peripheral blood of a 22-year-old man with CML in myeloid blast crisis (t(9;22)(q34;q11), e13a2 BCR-ABL). MV4-11 cells are macrophage type that stem from a 10-year-old boy with biphenotypic acute myelomonocytic leukemia and carry FLT3-ITD as oncogenic driver (kind gifts from Prof. Dr. Frank Dietmar Böhmer, University Clinic Jena, Germany). U937 and THP-1 cells are from the pleural effusion of a 37-year-old male patient with histiocytic lymphoma and from a 1-year-old boy with acute monocytic leukemia, respectively (both are kind gifts from Prof. Dr. Thomas Kindler (University Medical Center, Mainz, Germany). The cells were authenticated by DNA fingerprint profiling using eight different and highly polymorphic short tandems repeats, at the Leibniz-Institute DSMZ, Braunschweig, Germany). Leukemic cells were maintained in RPMI-1640 medium containing L-glutamine supplemented with 10% fetal bovine serum, 10 U/mL penicillin, and 10 µg/mL streptomycin. All cell lines were repeatedly tested for mycoplasma via PCR or enzymatic assays. We used only mycoplasma-free cells.

### Primaries and ethical approval

Patient-derived mononuclear cells from CML patients were collected upon approval of the Ethical Committee of Landesärztekammer Rhineland-Palatine No.: 837.258.17 (11092). were isolated by Ficoll Hypaque gradient and cultivated in L-glutamine containing Iscove’s Modified Dulbecco’s Medium (IMDM) (Thermo Fisher Scientific, Darmstadt, Germany) supplemented with 20% fetal bovine serum, 1% penicillin/streptomycin, 1x MEM non-essential amino acids (EuroClone), 10 ng/mL Interleukin-3, and 50 ng/mL FLT3 ligand (PeproTech/Thermo Fisher Scientific, Darmstadt, Germany). All cells were cultivated under sterile conditions at 37°C in a humidified incubator with 5% CO_2_.

### Inhibitors and drugs

MS-275, RGFP966, FK228, and imatinib mesylate were purchased from Selleck Chemicals. HU and cytarabine were from Sigma-Aldrich. Z-VAD-FMK was obtained from Bachem, MERCK60 (BRD6929) from Sigma. Marbostat-100 was synthesized and kindly provided by the working group of Prof. Siavosh Mahboobi in Regensburg, Germany. Z-VAD-FMK was applied for 1 h pre-treatment with other inhibitors. HU was always freshly prepared in PBS. Stock solutions of all other agents were prepared in DMSO and stored at -80°C.

### Cell lysis, SDS-PAGE and immunoblot

For lysate preparations leukemic cells were seeded into 60 mm dishes at a density of 2 × 10^5^ cells per mL. After 24 h, cells were treated with drugs for 24 and 48 h. Cells were harvested and cell pellets were lysed with 0.5% NET-N lysis buffer buffer (100 mM NaCl, 10 mM Tris-HCl pH 8, 1 mM EDTA, 10% glycerol, 0.5% NP-40; plus cOmplete protease inhibitor cocktail tablets (Roche, Mannheim, Germany) and phosphatase inhibitor cocktail 2 (Sigma-Aldrich, Germany)) containing DTT and phosphatase inhibitors. The Bradford-assay was carried out to determine protein concentrations of whole cell extracts. SDS-PAGE and immunoblotting were performed as recently described by us in (Fischer et al., 2023). Antibodies for immunoblot were purchased from Abcam: ab32420; BAK, ab32371; BAX, ab32503; FOXO1A, ab39670; HDAC3, ab32369; p-p73 (Tyr99), ab218625; p21, ab109199; α-tubulin, ab176560; BD Pharmingen™: BCL-XL, 51-6646GR; cleaved PARP1 (D214), 552596; PARP1, 556362; STAT5, 610192; Biozol: Vinculin, BZL-03106; Cell Signaling Technology: BCR, #3902; p-CHK1 (Ser317), #2344; CHK1, #2360; cleaved caspase-3 (Asp175), #9661; HDAC1, #34589; ɣH2AX (Ser139), #9718; histone H3, #9715; MYC, #5605; p73, #14620; Calbiochem: NOXA, OP180; Enzo Life Sciences: HSP90 (AC88), ADI-SPA-830-F; Novus Biologicals: survivin, NB500-201; Santa Cruz Biotechnology: β-actin, sc-47778; BCL2, sc-492; p-ABL (Tyr412), sc-293130; EGR1, sc-110; HDAC2, sc-7899; MCL1, sc-819; NF-кB p65, sc-372; TBP, sc-374035; Sigma-Aldrich: BIM, B7929; Thermo Fisher Scientific: ABL, ZC015; p-STAT5 (Tyr694), MA5-14973. Proteins were detected using enhanced chemiluminescence (Western Lightning Plus-ECL, Perkin Elmer, Rodgau, Germany) and the iBright CL1000 imaging system (Invitrogen, Thermo Fisher Scientific, Darmstadt, Germany) after incubation with anti-rabbit IgG HRP-linked/7074), anti-mouse IgG HRP-linked/7076 (Cell Signaling Technology, Frankfurt, Germany) or the infrared imager Odyssey (LICOR, NE, USA) after incubation with fluorescence-coupled LI-COR secondary antibodies (1:10,000) (IRDye^©^680RD anti-mouse IgG, # 925–68070; IRDye^©^680RD anti-rabbit IgG, #925–68071; IRDye^©^800CW anti-mouse IgG, #925–32210; IRDye^©^800CW anti-rabbit IgG, #925–32211). Densitometric evaluations were done with Image Studio Lite 4.0 (LI-COR, NE, USA). The untreated control was defined as 1.0 and measured intensities were normalized to the loading control.

### Biochemical cell fractionation

To prepare cytosolic and nuclear extracts, 1.5 × 10^5^ K562 cells per mL were seeded into 100 mm dishes. After 24 h, cells were treated with 5 µM MS-275 and 1 mM HU for 24 h. Cells were harvested into 15 mL falcons and centrifuged at 300 × g for 5 min at room temperature. The supernatant was discarded. The cells were then washed with 1 mL PBS and centrifuged again as stated before. All following steps were performed with pipettes having their tips cut to maintain the integrity of cell nuclei. Cell pellets were lysed in 400 µL hypotonic buffer (20 mM HEPES, pH 7.9, 10% glycerol, 0.2% NP-40, 10 mM KCL, 1 mM EDTA, 1 mM DTT, 0.5 mM PMSF, 1 tablet protease inhibitor cocktail (Roche, Mannheim, Germany) per 10 mL) for 7 min on ice. Nuclei were spun down for 2 min at 12,000 × g at 4°C. The supernatant containing the cytosolic fraction was collected in a fresh 1.5 mL tube. Protein concentration was determined by Bradford-assay. The nuclei were resuspended in 100 µL ROTI^®^Load Lämmli buffer (Roth, Karlsruhe, Germany) and heated for 5 min at 95°C. Afterwards, samples were analyzed by SDS-PAGE and immunoblotting.

### Cell cycle analysis

Leukemic cells were seeded into 12 well plates at a density of 2 × 10^5^ cells per mL. After treatment, cells were harvested, fixed with 80% ethanol, stained with the DNA intercalator dye propidium iodide (PI; Sigma-Aldrich, Steinheim, Germany) as described by us (Fischer et al., 2023; Mustafa et al., 2025), and measured on a FACS Canto II flow cytometer (BD-Biosciences). PI measurement revealing cell cycle distribution and DNA fragmentation (subG1 fraction) was performed. The obtained data were evaluated with the FACSDiva software 7.0 (BD-Biosciences).

### Analyses of apoptosis and mitochondrial membrane potential (ψm)

To distinguish apoptotic, necrotic, or living cells quantitatively, we used the fluorescent dyes FITC-labeled annexin-V (MACS Miltenyi Biotec) and PI (Sigma-Aldrich, Steinheim, Germany). Annexin-V-FITC is a marker for apoptosis, as it binds to the phospholipid phosphatidylserine which flips from the inner leaflet of the plasma membrane to the outer leaflet in apoptotic cells. It is possible to distinguish viable from nonviable necrotic cells with PI, being not permeable through intact membranes. Cells that are positive for annexin-V are early apoptotic and cells with an additional accumulation of PI are in late apoptosis (Beyer et al., 2017). Furthermore, cells were seeded into 12well plates at a density of 2 × 10^5^ cells per mL. For the DiOC6 staining to analyze the mitochondrial membrane potential (ψm) in cells, we used DiOC6(3) (Molecular Probes, Eugene). DiOC6(3) accumulates in the mitochondrial matrix upon changes in ψm (Δψm). DiOC6 staining was performed as previously described by us (Pons et al., 2018).

### RNA interference and plasmid transfection

The Amaxa® Cell Line Nucleofector^®^ Kit V (Lonza, Cologne, Germany)/Ingenio Solution Kit (Mirus; 100 µL solution per sample) was used for siRNA or plasmid transfections into suspension cells. Per sample, 1 × 10^6^ K562 cells were electroporated using The Amaxa^®^ Cell Line Nucleofector^®^ II device according to the protocol for K562 cells (ATCC^®^, VA, USA). We applied 100 pmol siRNA (nontargeting control siRNA-B/sc-44230 Santa Cruz; siRNA targeting NOXA/sc-37305 (Santa Cruz Biotechnology, Heidelberg, Germany); siRNA targeting p73/J-003331-10-0002 (Dharmacon, CO, USA); siRNA targeting HDAC1/L-003493-00-0005 (Dharmacon, CO, USA); siRNA targeting HDAC2/sc-29345 (Santa Cruz Biotechnology, Heidelberg, Germany); siRNA targeting HDAC3/sc-35538 (Santa Cruz Biotechnology, Heidelberg, Germany)) or 4 µg plasmid DNA. Cells were transferred to 6 well plates (1 × 10^6^ cells per well) in 4 mL fresh medium and incubated for 24 h. Afterwards, the cells were treated with drugs for 24-48 h.

### Generation of CRISPR-Cas9 knockout cells

Single guide RNAs (sgRNAs) were designed using the online tool Integrated Technologies (IDT), retrieved from https://www.idtdna.com. The most efficient guide RNA combinations were chosen based on the scoring system that is described in (Doffo et al., 2022). The sgRNAs were tested in pairs and the most efficient combination was kept for further knockouts. Exon 1 and exon 2 of the human *TP73* gene were targeted using the two sgRNAs 5′-*CACCTTCGACACCATGTCGC*-3′/5′-*GGTGCCCTATGAGCCACCACAG*-3′. The NOXA encoding gene *PMAIP1* was targeted using the two sgRNA sequences 5′-*TCGAGTGTGCTACTCAACTC*-3′/5′-*GCTACTCAACTCAGGAGATT*-3′. Ribonucleotide protein complexes of the sgRNAs and the GFP-Cas9 enzyme were electroporated into target cells using the Neon transfection system (Thermo Fisher Scientific). Transfection efficiency was assessed 24 h later by measuring the GFP signal with flow cytometry. GFP-positive cells were sorted and cultured as single cells in 96 well plates using the BD FACSAria™ III system. Single cells were maintained for approximately three weeks. Resulting single-cell colonies were screened for knockout efficiency by immunoblot.

### Preparation of RNA and quantitative real-time PCR

K562 cells were seeded at a density of 2 × 10^5^ cells per mL. Cells were treated with the indicated drugs for 24 h. Cell pellets from independent experiments were frozen at -80°C until RNA was isolated. Total RNA was isolated using the Nucleo Spin RNA Kit (Macherey and Nagel, Düren, Germany). Afterwards, 0.5 µg total RNA was transcribed into cDNA (Verso cDNA Kit, Thermo Scientific, Brunswick, Germany) and real time PCR was performed using the SensiMix^TM^SYBR Green & Fluorescein Kit (Bioline, London, UK) and the CFX96 Real-Time PCR Detection System (BioRad, Munich, Germany). PCR analyses were performed in technical triplicate, with SD showing intra-experimental variation. Analysis was performed using the CFX Manager^TM^ software. No-template controls were included in each run and expression levels of the genes of interest were normalized to *GAPDH* and *ACTB*; untreated control samples were set to one. The specific primers are depicted in **Table 1**.

**Table 1:**
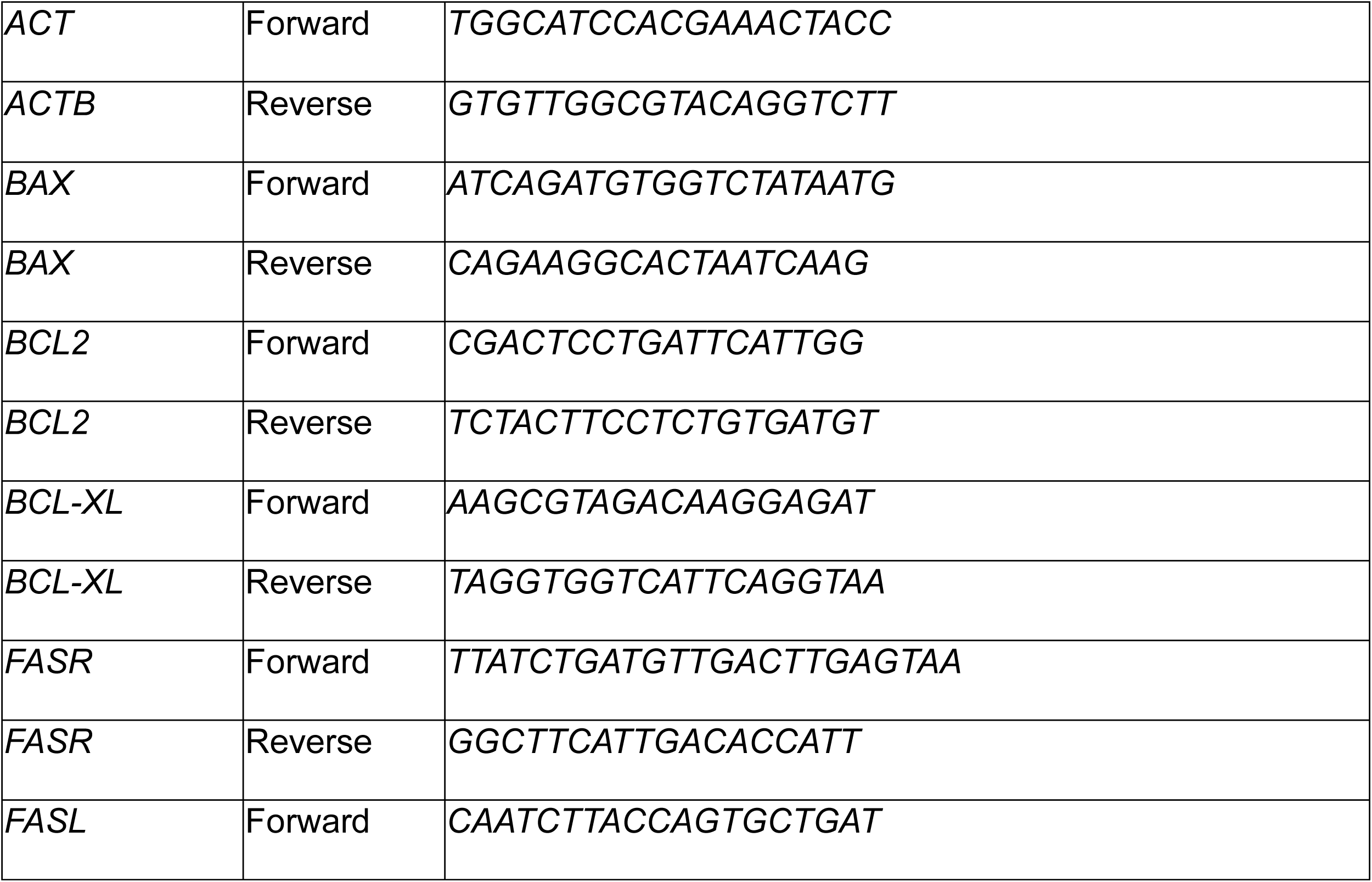
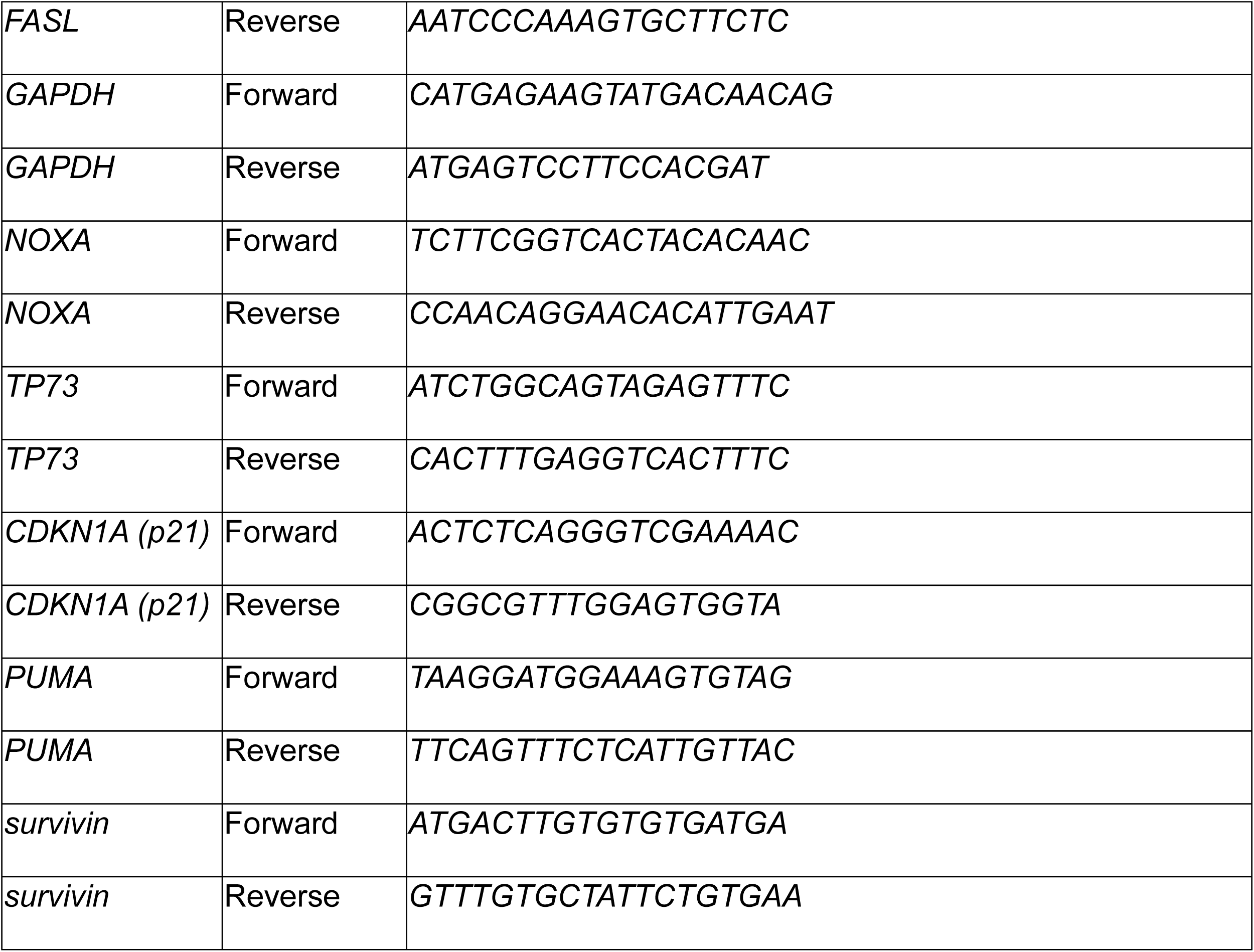
Primers

### Analysis of ENCODE datasets

Publicly available datasets from the ENCODE Project (Consortium, 2012) were utilized to investigate chromatin and transcriptional features associated with the *NOXA* gene in K562 cells. We analyzed ChIP-sequencing data for the histone modification H3K27ac (ENCODE ID: ENCSR000AKP), the insulator CTCF (ENCODE ID: ENCSR000DWE), and the histone deacetylases HDAC1 (ENCODE ID: ENCSR711VWL), HDAC2 (ENCODE ID: ENCSR893WSB), HDAC3 (ENCSR024LKA), HDAC6 (ENCSR000ATJ), and HDAC8 (ENCSR835TCD) to assess chromatin binding profiles. In parallel, ATAC-sequencing data (ENCODE ID: ENCSR868FGK) were examined to evaluate chromatin accessibility. RNA-sequencing data (ENCODE ID: ENCLB555AML) were analyzed to assess gene expression. All datasets were downloaded from the ENCODE data portal (https://www.encodeproject.org) and processed using the Integrated Genome Viewer (IGV) (Robinson et al., 2011), a tool for visualization and analysis of the *NOXA* gene region across datasets. ChIP-sequencing peaks, ATAC-sequencing signals, and RNA-sequencing reads were aligned to the human genome (hg38) using similar peak intensity settings in IGV. Regions of interest were defined by peaks overlapping the promoter and enhancer regions of the *NOXA* gene. These data were integrated to evaluate CTCF and HDAC occupancy, histone acetylation, chromatin accessibility, and gene expression of *NOXA*.

### Analysis of the database HEMAP

The publicly available hematopoietic database HEMAP is a curated compendium of gene expression patterns in normal blood cells, hematologic malignancies, and leukemia subtypes. Data contained in this data collection were obtained by transcriptomics (Pölönen et al., 2019).

## Statistical analyses

Depending on the experimental setup, t-test, one-way or two-way ANOVA (Bonferroni’s multiple comparisons test) were applied to determine statistical significance. p-values were defined as: p*<0.05, p**<0.01, p***<0.001, p****<0.0001. The tests were performed with the GraphPad Prism 6 software. Unless otherwise stated, all experiments were performed in biological triplicates. The specifics of the statistical methods used are described in figure legends.

## Results

### Class I HDACs are required for the survival of CML cells with DNA replication stress

We previously demonstrated that HU stalled CML cells in the S phase of the cell cycle without inducing apoptosis (Pons et al., 2018; Pons et al., 2021b). This characteristic resistance of CML cells to HU-induced apoptosis makes HU-treated CML cells an ideal model to investigate DNA replication independently of cell death-associated processes. Due to their increasingly appreciated roles as stress regulators, we speculated that HDACs control apoptosis induction upon HU-induced DNA replication stress. To probe the role of HDAC1, HDAC2, and HDAC3 in this context, we used entinostat (MS-275). This benzamide-based HDACi targets these HDACs specifically (Bradner et al., 2010). We treated K562 cells, which carry BCR-ABL, with 5 µM MS-275 and 1 mM HU for 24-48 h. With lysates from them we estimated the activation of the ultimate apoptosis executioner enzyme caspase-3 by immunoblot (Marx-Blümel et al., 2017). This analysis revealed robust activation of caspase-3 in MS-275-treated K562 cells. Although HU did not activate caspase-3, it significantly enhanced caspase-3 activation by MS-275 (**Fig. 1A**). Cleavage of PARP1 into an 89 kDa fragment is a hallmark of apoptotic execution as it verifies the activation of caspase-3 against one of its target proteins (Marx-Blümel et al., 2017). We could corroborate caspase-3 activation as accumulation of cleaved PARP1 (**Fig. 1A**). The key molecular marker for class I HDAC inhibition is an accumulation of acetylated histones (Beyer et al., 2017). Immunoblot analyses showed that these pro-apoptotic effects were not attributable to changes in the hyperacetylation levels of histone H3 (**Fig. 1A**). These findings suggest that a specific regulatory mechanism can explain apoptosis induction by HU plus MS-275.

**Fig. 1:**
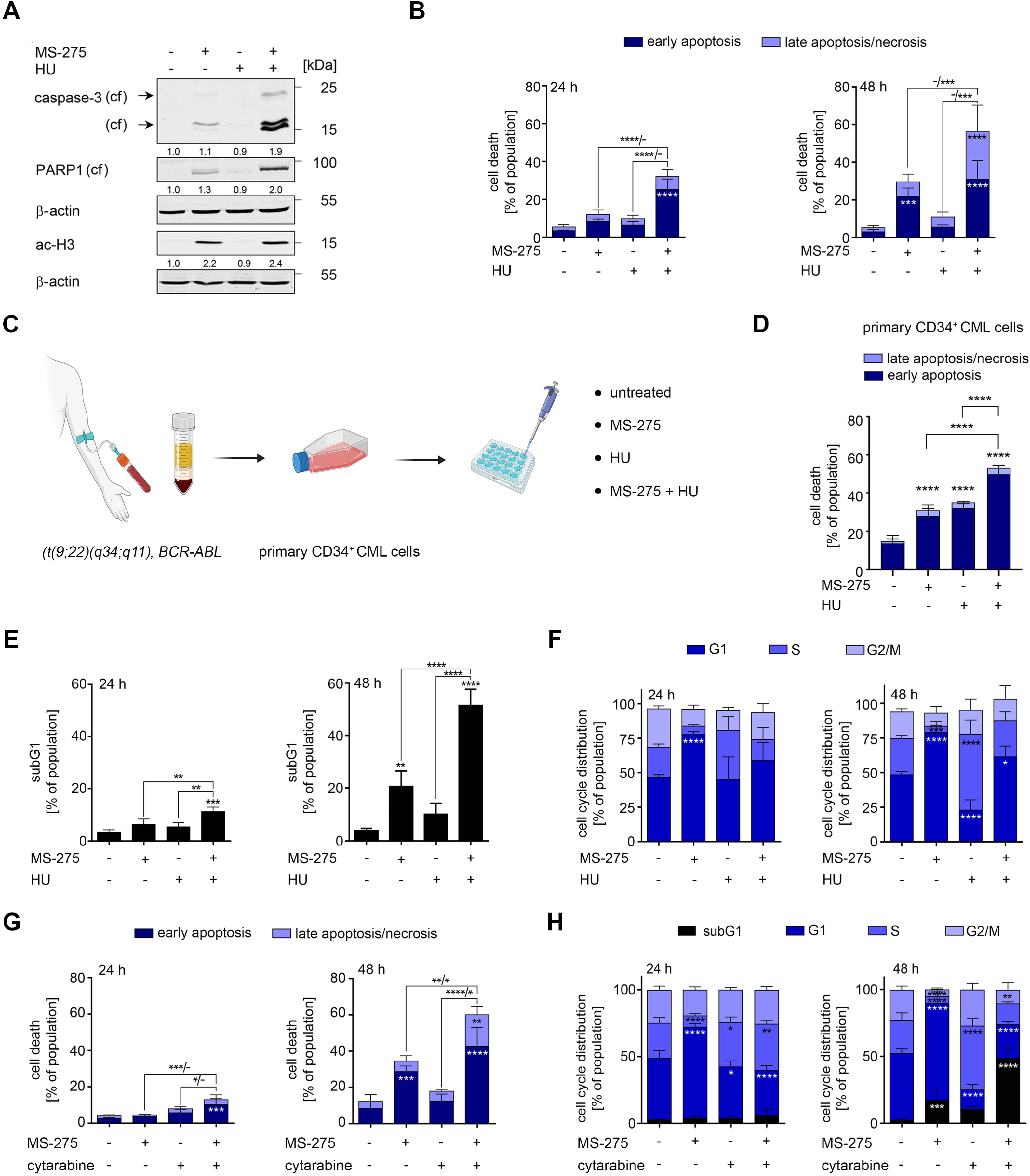
Inhibition of class I HDACs induces apoptosis in K562 cells undergoing DNA replication stress. A) K562 cells were treated with 1 mM HU ± 5 µM MS-275 for 24 h. Cells were lysed and indicated proteins were analyzed by immunoblot (cf, cleaved forms, arrows; ac, acetylated). β-actin serves as loading control. Densitometric analyses display relative expression of the indicated proteins. Values were normalized to the loading controls of the respective membranes; n=3. B) K562 cells were treated with 1 mM HU ± 5 µM MS-275 for 24 h and 48 h. Flow cytometry was performed to measure annexin-V/PI-positive cells; n=4, mean±SD; two-way ANOVA, Bonferroni’s multiple comparisons test; p***<0.001, p****<0.0001. C) Scheme for the collection of CML cell primaries and for their treatment ex vivo. Whole blood is collected from the arm crook, white blood cells are isolated by centrifugation, cultured, and treated as indicated. In the acute phase of CML, most white blood cells are BCR-ABL-positive CML cells. D) Primary CD34^+^ CML cells were treated with 1 mM HU ± 5 µM MS-275 for 48 h. Flow cytometry was performed to measure annexin-V/PI-positive cells; n=3, mean±SD; two-way ANOVA, Bonferroni’s multiple comparisons test; p***<0.001, p****<0.0001. E) SubG1 fractions in PI-stained K562 cell cultures were determined by flow cytometry; treatment conditions are given in A); n=4, mean±SD; one-way ANOVA, Bonferroni’s multiple comparisons test; p**<0.01, p***<0.001, p****<0.0001. F) K562 cells treated as mentioned in A), and cell cycle distributions were determined of PI-stained cells by flow cytometry; n=4, mean±SD; two-way ANOVA, Bonferroni’s multiple comparisons test; p*<0.05, p***<0.001, p****<0.0001. G) K562 cells were treated with 5 nM MS-275 ± 1 µM cytarabine for 24 and 48 h. Flow cytometric analysis was done to measure annexin-V/PI-stained cells; n=3, mean±SD; ANOVA, Bonferroni’s multiple comparisons test n; p*<0.05, p**<0.01, p***<0.001, p****<0.0001. H) K562 cells were treated as indicated in H) and stained with PI for cell cycle distribution analysis; n=3, mean±SD; ANOVA, Bonferroni’s multiple comparisons test; p*<0.05, p**<0.01, p***<0.001, p****<0.0001.

Next, we employed flow cytometry for cell surface-exposed annexin-V and propidium iodine (PI) to quantify the levels of apoptosis induction by MS-275±HU. HU did not induce apoptosis significantly after 24 h and 48 h. MS-275 caused apoptosis of K562 cells significantly after 48 h (**Fig. 1B**). The combinatorial treatment of HU and MS-275 provoked apoptosis significantly after 24 h and 48 h (**Fig. 1B** and **S1A**). Moreover, the combination of HU and MS-275 was significantly more pro-apoptotic than the single agents and particularly induced the irreversible state of late apoptosis after 48 h (**Fig. 1B** and **S1A**). To investigate these data in a primary patient-derived CML model, we studied if MS-275 induced apoptosis in freshly isolated cells from a 39-year-old female with chronic phase t(9;22)-positive CML carrying BCR-ABL b2a2. We ensured their purity as positivity for the surface stem cell marker CD34^+^ (**Fig. 1C**). Flow cytometry demonstrated that HU and MS-275 induced apoptosis significantly in patient-derived leukemic blasts after 48 h. Compared to both single agents, the combination of HU and MS-275 was more effective and caused up to 53% apoptosis ex vivo (**Fig. 1D**).

To obtain a more complete apoptosis profile, we determined the occurrence of a subG1 phase. This fraction of cells confirms irreversible DNA fragmentation during the execution phase of late apoptosis (Marx-Blümel et al., 2017). Consistent with **Figs. 1B** and **S1A**, we observed that MS-275 significantly and dose-dependently increased the subG1 fraction after 48 h. HU plus MS-275 triggered apoptotic DNA fragmentation at 24 h and 48 h significantly and more potently than single treatments (**Fig. 1E**).

To account for potential cell line-specific effects, we analyzed the impact of HU and MS-275 treatment on apoptosis induction in additional BCR-ABL-positive human CML cell types. We used LAMA-84, MEG-01, KCL-22, and KYO-01 cells. Like in K562 cells, combined application of HU and MS-275 evoked apoptotic DNA fragmentation more strongly than the individual drugs in all four cell lines (**Fig. S1B**). These data verify that CML cells share susceptibility to MS-275 and that the inhibition of HDAC1/2/3 by MS-275 triggers apoptosis in various CML systems undergoing DNA replication stress.

Next, we analyzed how HU and MS-275 altered cell cycle progression by flow cytometry. We incubated K562 cells with HU ± MS-275 for 24-48 h. MS-275 caused a significant G1-phase arrest, which lasted for 48 h, and subsequently depleted the cell populations in the S and G2/M phases (**Fig. 1F** and **S1C**). Consistent with our previous data for HU-treated K562 cells (Pons et al., 2019; Pons et al., 2018; Pons et al., 2021a), we noted that HU stalled the cell cycle in the S phase at 24 h. This effect became more pronounced and highly significant at 48 h (**Fig. 1F** and **S1C**). The cotreatment evoked intermediate effects, with a detectable but not significant dominance of MS-275 effect at 24 h. After 48 h, a significant portion of the remaining vital K562 cells in the cotreatment group were in G1 phase, suggesting that MS-275 halts DNA replication. The addition of MS-275 to HU-treated cells caused a depletion of the S phase cell population after 48 h. Although most K562 cells that were plated with MS-275 for 48 h were in G1 phase, the addition of HU attenuated this effect (**Fig. 1F** and **S1C**). These findings show that the biological responses of cells to single drugs are partially abolished upon their combination.

We aimed to substantiate the above findings for HU with another DNA replication disrupter. The antimetabolite cytarabine stalls cells in S phase and is a frequently used medication for leukemia (Gielecinska et al., 2023; Park et al., 2020). We noted that K562 cells did not undergo apoptosis upon treatment with cytarabine for up to 48 h. The combined application of cytarabine and MS-275 induced early and late apoptosis significantly at 24 h and 48 h, reaching 58% of total apoptosis (**Fig. 1G**). Evaluation of the cell cycle and subG1 phase showed that cytarabine caused an accumulation of K562 cells in S phase after 24 h. The portion of S phase cells increased after a 48-h treatment with cytarabine. Although MS-275 did not halt the cytarabine-induced arrest in G1 phase after 24 h, MS-275 reduced the percentage of cytarabine-treated K562 cells in S phase. After 48 h, cytarabine plus MS-275 caused apoptotic DNA fragmentation and ceased the cytarabine-induce S phase arrest significantly (**Fig. 1H**). These data show that our data for HU can be recapitulated with cytarabine.

We additionally assessed if other HDACi combine favorably with HU. The depsipeptide FK228, which inhibits the four class I deacetylases HDAC1,-2,-3,-8 (Mustafa and Krämer, 2023), significantly augmented apoptosis and DNA fragmentation in HU-treated K562 cells after 24 h and 48 h (**Figure S2A,B**). Like with MS-275, such cell death was linked to a depletion of K562 cells in S phase (**Figure S2A,B**). Since the class IIB deacetylase HDAC6 is frequently dysregulated in leukemic cells, we treated K562 with the highly selective HDAC6 inhibitor marbostat-100 (Beyer et al., 2019; Sellmer et al., 2018). Unlike MS-275 and FK228, marbostat-100 did not activate apoptosis or DNA fragmentation in HU-treated K562 (**Figure S2C,D**).

These data demonstrate that inhibitors of HDAC1, HDAC2, and HDAC3 sensitize cultured and primary CML cells to apoptosis induction upon DNA replication stress induction.

### MS-275 and HU upregulate NOXA expression in CML cells

To identify the molecules that regulate apoptosis induction by HU plus MS-275 in K562 cells, we carried out a small-scale microarray for the expression of pro-apoptotic and anti-apoptotic proteins. As positive control for the effectiveness of MS-275 and HU, we analyzed the mRNA for the cell cycle regulator *p21* which is induced by both agents (Göder et al., 2018). We noted a significant upregulation of the mRNA encoding the pro-apoptotic BH3-only protein NOXA by MS-275 and HU in combination but not of the other pro-apoptotic molecules; *p21* accumulated as expected (**Fig. 2A**).

**Fig. 2:**
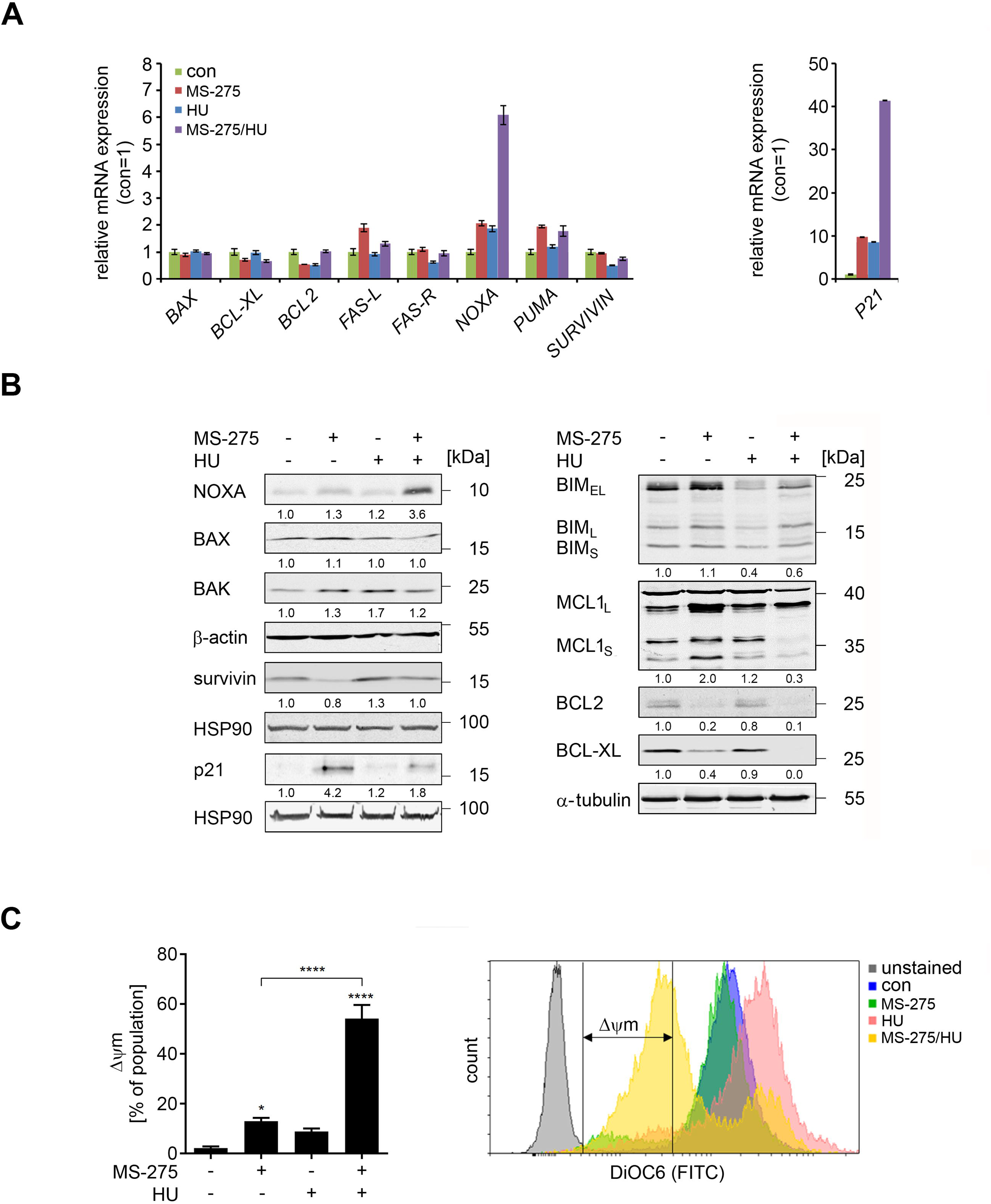
K562 cells treated with HU and MS-275 accumulate NOXA mRNA and protein. A) K562 cells were treated with 1 mM HU ± 5 µM MS-275 for 24 h. A qRT-PCR screen revealed gene expression signatures as shown. Expression was normalized to *GAPDH* and *ACTB*. B) K562 cells were treated with 1 mM HU ± 5 µM MS-275 for 24 h and 48 h. Cells were lysed and analyzed for the indicated proteins by immunoblot; β-actin and HSP90 serve as loading controls. The numbers below the blots show densitometric evaluation of relative protein expression. Values were normalized to loading controls. C) K562 cells were stained with DiOC6 and analyzed by flow cytometry to determine changes in the mitochondrial membrane potential (Δ*Ψ*m); n=3 ± SD; one-way ANOVA; Bonferroni’s multiple comparisons test; p*<0.05, p****<0.0001.

To substantiate these data, we analyzed the expression of the proteins that are encoded by these mRNAs and of further apoptosis regulators by immunoblot. Consistent with the mRNA expression data for *NOXA* (**Fig. 2A**), we detected an accumulation of NOXA (**Fig. 2B**). HU and MS-275 did not or weakly alter the expression of BAX and BAK, which can both form mitochondrial pores that allow the efflux of cytochrome-c and subsequent caspase activation (Shamas-Din et al., 2011). Unlike in solid tumor-derived cells (Stauber et al., 2012), MS-275 and HU did not increase the levels of pro-apoptotic BIM (**Fig. 2A**). We further found that MS-275 increased the small and large isoforms of MCL1, whereas HU and HU/MS-275 reduced MCL1. MS-275 reduced BCL-2 and BCL-XL levels and this effect was further accentuated when HU was added to MS-275. MS-275 attenuated the levels of survivin, but HU prevented this (**Fig. 2B**). Unlike the accumulation of *p21* mRNA by HU ± MS-275 (**Fig. 2A**), p21 did only accumulate in MS-275-treated K562 cells (**Fig. 2B**). This apparent discrepancy can be explained by an inhibition of *p21* mRNA translation by HU (Beckerman et al., 2009). The reduction of p21 in MS-275-treated K562 cells by HU corresponds to a decrease in G1 phase cells when HU is given with MS-275 (**Fig. 1F and S1C**).

These results demonstrate that HU and MS-275 alter the levels of pro- and anti-apoptotic factors that are responsible for maintaining mitochondrial integrity. To evaluate this, we analyzed the loss of the mitochondrial transmembrane potential (Δψm), being an early sign of apoptosis. We noted a significant mitochondrial injury in 55% of K562 cells that were treated with HU and MS-275 for 24 h (**Fig. 2C**).

These data disclose that HU and MS-275 induce NOXA and reduce anti-apoptotic BCL2 family members.

### Apoptosis induction by MS-275 and HU in CML cells requires NOXA

To determine if the upregulation of NOXA following combined treatment with HU and MS-275 has functional relevance for the induction of apoptosis, we applied two complementary genetic approaches. As a first strategy, we depleted NOXA expression using RNAi. Immunoblot analyses for PARP1 cleavage revealed that treatment with 1.5–5 µM MS-275 in combination with 1 mM HU markedly increased the levels of cleaved PARP1 in K562 cells that we transfected with control siRNAs. Notably, this accumulation of cleaved PARP1 was completely prevented upon NOXA depletion by RNAi (**Fig. 3A**). We controlled this experiment by immunoblot for phosphorylated H2AX (ɣH2AX) which accumulates upon HU-induced DNA replication stress and damage (Göder et al., 2018). The genetic depletion of NOXA by its cognate siRNAs did not affect the induction of ɣH2AX by HU ± MS-275 in K562 cells (**Fig. 3A**). Measurement of early and late apoptosis by flow cytometry confirmed the data that we collected for cleaved PARP1. The knockdown of NOXA by siRNA significantly attenuated apoptosis induction following treatment with 1 mM HU and 1.5-5 µM MS-275 for 24 h (**Fig. 3B**).

**Fig. 3:**
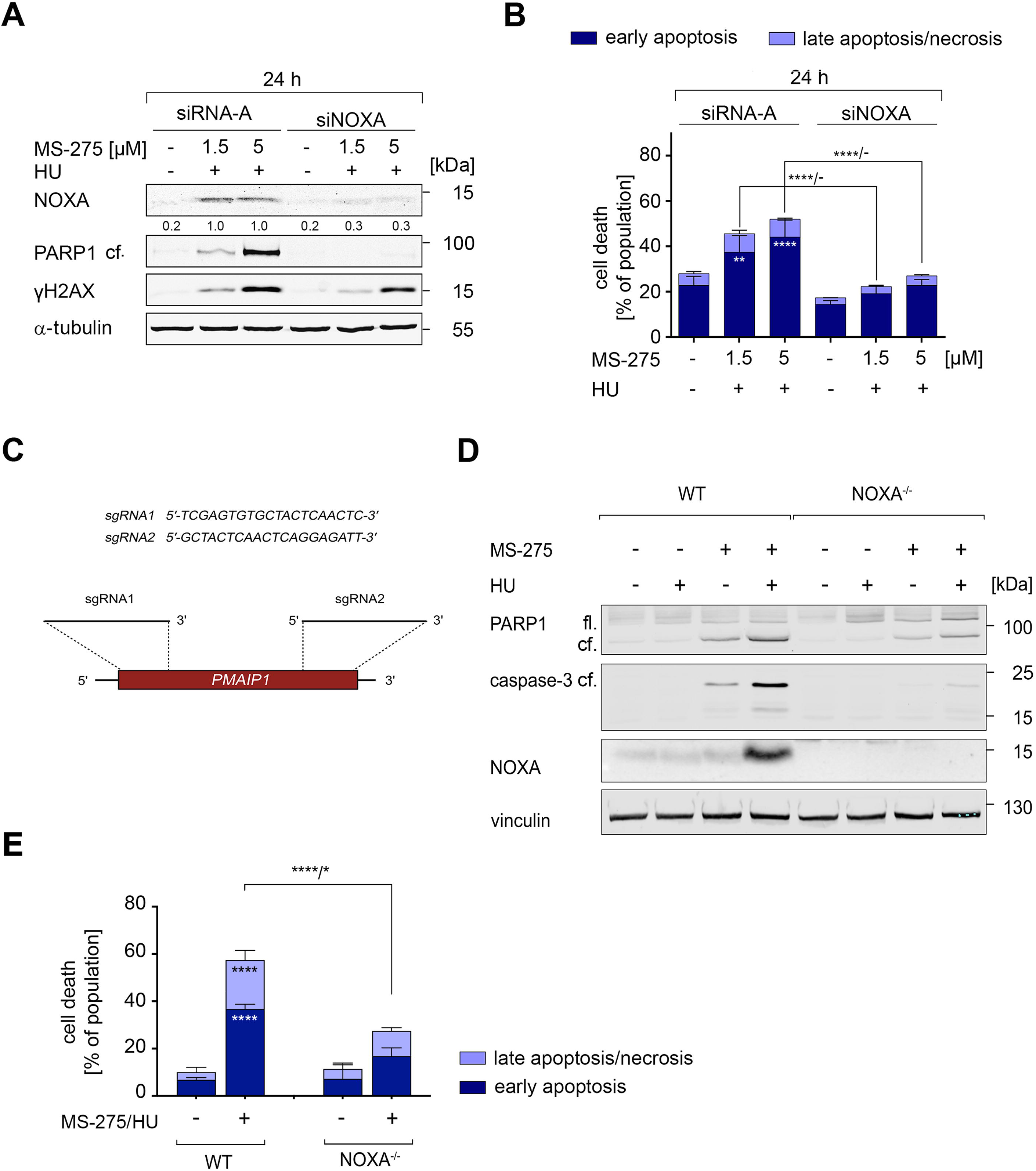
NOXA is required for apoptosis in cells undergoing DNA replication stress. A) Knockdown of NOXA in K562 cells with RNAi. Following electroporation, cells were treated with 1.5-5 µM MS-275 ± 1 mM HU for 24 h, analyzed for NOXA, γH2AX and the cleavage of PARP1 by immunoblot; α-tubulin and β-actin serve as loading control. The numbers below the blots indicate densitometric evaluation of relative protein expression. Values were normalized to the loading control. B) Cells were treated as in (A), stained with annexin-V/PI, and subjected to flow cytometry; n=3 ± SD; two-way ANOVA, Bonferroni’s multiple comparisons test; p*<0.05, p**<0.01, p****<0.0001. C) Schematic diagram showing genetic knockout of NOXA encoding gene *PMAIP1* using the indicated sgRNAs by CRISPR-Cas9 technology. Wild-type and *NOXA* ^-/-^ K562 were treated with 5 µM MS-275 ± 0.5 mM HU for 48 h, stained with annexin-V/PI, and subjected to flow cytometry; n=3 ± SD; two-way ANOVA, Bonferroni’s multiple comparisons test; p*<0.05, p**<0.01, p****<0.0001. D) Wild-type (WT) and *NOXA* null (NOXA^-/-^) K562 cells were treated as stated in C). The cells were analyzed for NOXA and the cleavage of caspase-3 and PARP1 by immunoblot; vinculin as loading control. The numbers below immunoblots are densitometric evaluation of relative protein expression. These are normalized values in relation to the loading control.

To corroborate these data with a complete genetic elimination of NOXA, we disrupted the NOXA encoding gene *PMAIP1* using the CRISPR-Cas9 technology (**Fig. 3C**). Immunoblotting verified that K562 cells lacking NOXA (K562^NOXA-/-^) did not carry NOXA and did not accumulate it upon treatment with MS-275, HU, and their combination (**Fig. 3D**). We could reproduce these data in three independent clones lacking NOXA (**Fig. S3A**) which rules out clonal effects. We used wild-type K562 cells (K562^WT^) as positive control for the induction of NOXA under these conditions (**Fig. 3D, S3A**).

We analyzed the activation of caspase-3 and the resulting cleavage of PARP1 in control K562 cells (K562^WT^) and in NOXA-negative K562^NOXA-/-^ cells that we plated with MS-275 and HU. Unlike seen in K562^WT^ cells, there was no induction of NOXA in K562^NOXA-/-^ cells (**Fig. 3D**). Consistent with this lack of biochemical pro-apoptotic signatures, flow cytometry illustrated that K562^NOXA-/-^ cells did not undergo apoptosis significantly when incubated with MS-275 and HU. Under equal conditions, K562^WT^ cell cultures had a significant induction of early and late apoptosis (**Fig. 3E**). Consistent with a lack of NOXA induction by HU plus MS-275 in several K562^NOXA-/-^ cell cultures (**Fig. S3A**), these did not do into apoptosis (**Fig. S3B**). Thus, clonal effects can be excluded.

These results demonstrate that NOXA is required to induce apoptosis in CML cells undergoing DNA replication stress and concomitant class I HDAC inhibition.

### BCR-ABL regulates p73 stabilization and the resulting NOXA expression upon DNA replication stress and HDAC inhibition

Having defined the pivotal role of NOXA in our experimental settings, we next sought to identify its upstream regulators. The expression of *NOXA* mRNA is modulated by several transcription factors including EGR1, E2F1, FOXO1, MYC, NF-кB p65, p53, p73, and SP1, under various experimental conditions (Bertin-Ciftci et al., 2013; Grande et al., 2012; Lin et al., 2012; O’Prey et al., 2010; Wirth et al., 2014). Whether DNA replication stress and HDACi culminate in NOXA expression via these transcription factors remains to be clarified. We found that MS-275 plus HU reduced the levels of EGR1, FOXO1, MYC, and NF-кB p65, and K562 cells lack p53 (**Fig. S4A**). Genetic manipulation of MYC using RNAi or MYC overexpression consistently showed no impact on cell death induced by MS-275 plus HU (**Fig. S4B**).

In contrast to these results that do not support a role for EGR1, FOXO1, MYC, and NF-кB p65, p53, and E2F1, we observed that HU induced an accumulation of Tyr99-phosphorylated (p-p73) and total p73α in K562 cells. Upon addition of MS-275, the HU-induced increase in p73 persisted and MS-275 augmented the HU-evoked accumulation of p-p73 (**Fig. 4A**). Data obtained by qPCR showed that HU plus MS-275 did not augment *TP73* mRNA levels (**Fig. S4C**), indicating p73 stabilization at the protein level. Consistent with this idea, the accumulation of p73 is linked to its phosphorylation at Tyr99 which stabilizes p73 (Cai et al., 2022; Omran et al., 2021).

**Fig. 4:**
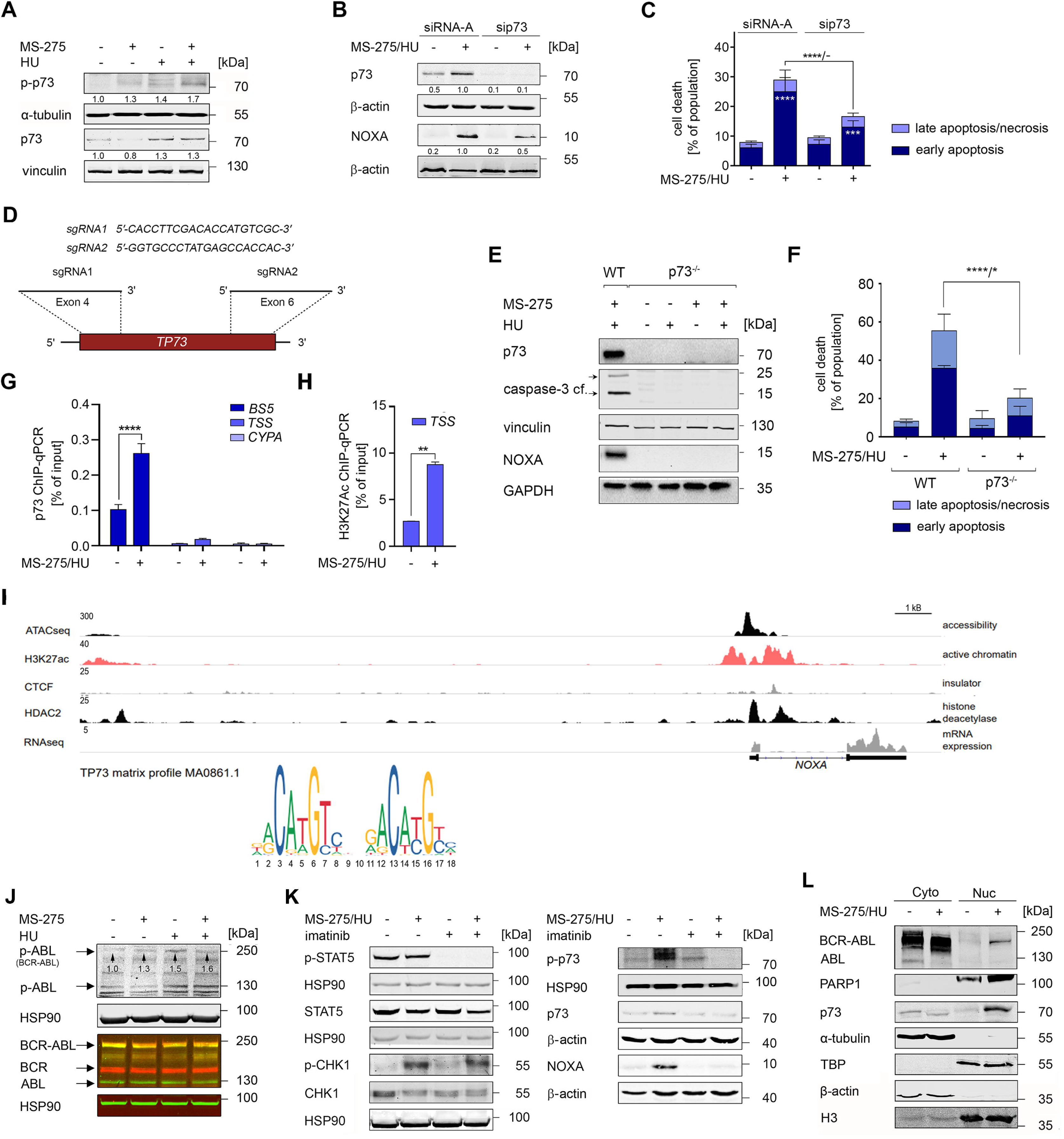
The transcription factor p73 regulates NOXA expression dependent on BCR-ABL in cells DNA replication stress. A) K562 cells were treated with 1 mM HU and 5 µM MS-275 for 24 h and analyzed for p-p73 (Tyr99) and p73 expression via immunoblot; vinculin serves as loading control. B) K562 cells were transfected with noncoding siRNAs or siRNAs against p73 and treated with 5 µM MS-275 and 1 mM HU for 24 h. Induction of the p73-NOXA signaling node by HU plus MS-275 and the knockdowns of p73 by RNAi were verified by immunoblot; β-actin serves as loading control. Numbers below are densitometric evaluations of relative protein expression. Values were normalized to corresponding controls. C) Percentages of annexin-V/PI positive cells in K562 cells that were transfected and treated as stated in B) were determined by flow cytometry; n=8 ± SD; two-way ANOVA; Bonferroni’s multiple comparisons test; p***<0.001, p****<0.0001. D) Schematic diagram showing genetic knockout of *TP73* using the indicated sgRNAs by CRISPR-Cas9 technology. E) Wild-type K562 cells (WT; positive control) and p73 knockout (p73^-/-^) K562 cells that were treated with 5 µM MS-275 + 0.5 mM HU for 24 h were analyzed for p73, NOXA and the cleavage of caspase-3 and PARP1 by immunoblot; vinculin and GAPDH serve as loading control. F) Wild-type (WT) and *p73* null (-/-) K562 cells were treated with 5 µM MS-275 + 0.5 mM HU for 48 h, stained with annexin-V/PI, and subjected to flow cytometry; n=3 ± SD; two-way ANOVA, Bonferroni’s multiple comparisons test; p*<0.05, p**<0.01, p****<0.0001. G) ChIP analysis of K562 cells treated with 1 mM HU and 5 µM MS-275 for 24 h shows binding of p73 to the *BS5* binding site in the *NOXA* promoter. ChIP was followed by qPCR; enrichment is given as % of input. The *cyclophilin* A (*CYPA)* promoter is used as negative control verifying specificity of the antibodies and the ChIP analysis; *transcriptional start site* (*TSS*). H) ChIP analysis performed as in F) also shows acetylation of the *NOXA* promoter at H3K27 in the *transcriptional start site* (*TSS*). I) ENCODE datasets were analyzed using ChIP-seq, ATAC-seq, and RNA-seq data in K562 cells to investigate chromatin binding profiles, accessibility, and transcriptional features of the *NOXA* gene. Data were processed for regions of interest, defined by peaks overlapping the *NOXA* promoter region. J) K562 cells were treated with 5 µM MS-275 ± 1 mM HU for 24 h. Lysates of the cells were analyzed for p-ABL (Tyr412) and BCR-ABL, BCR, and ABL by immunoblot; HSP90 serves as loading control. K) K562 cells were treated with 1 mM HU and 5 µM MS-275 ± 1 µM imatinib for 24 h. The indicated proteins were analyzed by immunoblot; HSP90 and β-actin are loading controls. The phosphorylation of p-STAT5 (Tyr694) was determined to verify BCR-ABL inhibition. L) K562 cells were treated with 5 µM MS-275 and 1 mM HU. Cell fractionation enabled the analysis of protein localization of BCR-ABL, c-ABL and p73; PARP1, α-tubulin, TBP, β-actin and histone 3 (H3) were used to show purity of cytoplasmic (Cyto) and nuclear (Nuc) fraction and equal loading.

Since the data above suggest a p73-dependent control of NOXA expression under our experimental conditions, we investigated the role of p73 on cell fate. Following our strategy for NOXA, we employed two different genetic strategies to deplete NOXA in K562 cells. An siRNA-mediated depletion of p73 significantly abolished the induction of NOXA and apoptosis induction in response to MS-275 plus HU (**Fig. 4B,C**). Encouraged by these data, we disrupted the *TP73* gene by CRISPR-Cas9 (**Fig. 4D**). Like K562^NOXA-/-^ cells (**Fig. 3D,E**), K562^p73-/-^ cells did not activate the p73-NOXA signaling node and did not accumulate cleaved caspase-3 when plated with MS-275±HU (**Fig. 4E**). Consistent herewith, K562^p73-/-^ cells did not become significantly annexin-V/PI-positive upon incubation with MS-275 plus HU (**Fig. 4F**). This lack of apoptosis induction in K562 cells lacking p73 was reproducible over various K562^p73-/-^ single-cell clones (**Fig. S3C,D**), disfavoring clonal effects.

To assess whether p73 directly activates NOXA expression in cells treated with MS-275 plus HU, we carried out quantitative chromatin immunoprecipitation (ChIP). We assumed that p73 binds to the identified proximal p53 consensus binding site in the *NOXA* gene promoter (*BS5*; (Grande et al., 2012)). We verified that p73 specifically binds to *BS5* upon the combination treatment by ChIP (**Fig. 4G**). This binding of p73 tied in with an accumulation of histone H3 acetylated at lysine 27 (H3K27ac) within the transcriptional start site (*TSS*; **Fig. 4H**).

We used ENCODE datasets (Consortium, 2012) to test these results and to gain insights into chromatin binding profiles, chromatin accessibility, and transcriptional features of *NOXA*. We studied ChIP-sequencing, ATAC-sequencing, and RNA-sequencing data that are available for *NOXA* in K562 cells. Here, we observed open chromatin at the proximal *NOXA* promoter harboring a p73 binding motif, together with active chromatin, and a detectable recruitment of HDAC2 (**Fig. 4I**).

To evaluate if the induction of p73 and NOXA upon inhibition of class I HDACs and DNA replication stress represents a general phenomenon of DNA replication stress, we treated K562 cells with MS-275 and cytarabine. The combined application cytarabine and MS-275 augmented the levels of p73 and NOXA significantly (**Fig. S4D**).

Since a combinatorial treatment of HU and MS-275 caused an accumulation of p73 and its phosphorylation of p73 at Tyr99, without changing its mRNA expression (**Fig. 4A** and **Fig. S4C**), we speculated that p73 was stabilized through posttranslational modifications. The DNA damage-induced ABL kinase stabilizes p73 (Cai et al., 2022; Omran et al., 2021). CML cells carry constitutively active BCR-ABL. To analyze their functions in our experimental settings, we first tested the levels of phosphorylated and total ABL as well as BCR-ABL in K562 cells treated with HU and MS-275. Alone and combined these agents neither affected the phosphorylated levels of BCR-ABL and ABL nor the total levels of BCR-ABL, ABL, and BCR significantly (**Fig. 5J**). We next investigated if the accumulation of NOXA upon treatment with HU and MS-275 for 24-48 h occurred in leukemia cells without BCR-ABL. In MV4-11, THP1, U937, and HEL cells, these agents did not induce NOXA (**Fig. S5A,B**).

**Fig. 5:**
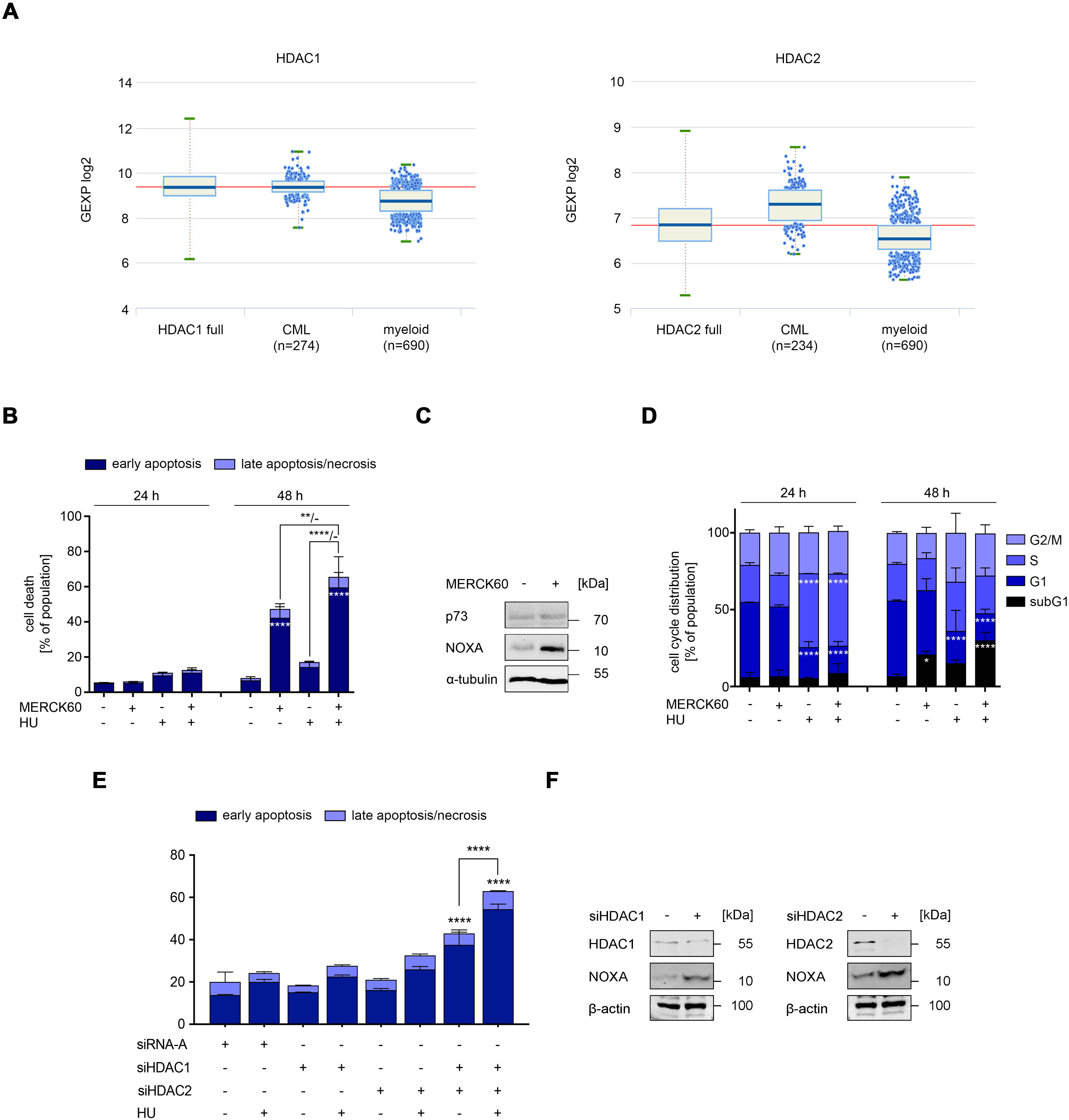
HDAC1 and HDAC2 regulate NOXA expression and apoptosis upon DNA replication stress. A) The Hemap database (http://hemap.uta.fi/) shows the mRNA expression levels of HDAC1 (n=274) and HDAC2 (n=234) in human CML cell samples compared to normal myeloid cells (n=690); full represents the maximum variation levels of these HDACs across all leukemia cell types and normal blood cells. B) K562 cells were treated with 1 mM HU ± 5 µM MERCK60 for 24 and 48 h. Annexin-V/PI-positive cell counts were measured by flow cytometry; n=3, mean±SD two-way ANOVA, Bonferroni’s multiple comparisons test; p**<0.01, p****<0.0001. C) Lysates were prepared from cells that were plated as mentioned in B) to detect p73 and NOXA protein; α-tubulin, loading control. D) K562 cells were treated as noted in B) and analyzed for cell cycle distribution via flow cytometry; n=3, mean±SD; two-way ANOVA, Bonferroni’s multiple comparisons test; p**<0.05, p****<0.0001. E) K562 cells were transfected with non-coding siRNA (siCon), siRNA against HDAC1 or siRNA against HDAC2. After 24 h, the cells were treated with 1 mM HU for 48 h. Annexin-V/PI-stained cell counts were measured via flow cytometry; n=3, mean±SD. F) Immunoblot shows HDAC1, HDAC2, and NOXA expression in K562 cells that were electroporated with siCon, siHDAC1, or siHDAC2; β-actin serves as loading control.

We used the clinically established tyrosine kinase inhibitor imatinib (Hochhaus et al., 2020) to test the functionality of the constitutive active BCR-ABL kinase for the p73-NOXA axis. Using the BCR-ABL target protein STAT5 (Ikeda et al., 2022), we identified that 1 µM imatinib blocked BCR-ABL activity in untreated and stressed K562 cells completely (**Fig. 5K**). This allowed us to scrutinize if BCR-ABL is mechanistically linked to increased levels of p-p73, total p73, and NOXA in these cells. The breakdown of BCR-ABL signaling by imatinib reduced phosphorylated p73 and abrogated the p73-NOXA signaling in K562 cells significantly (**Fig. 5K**). These observations cannot be explained by generally altered DNA replication stress signaling, which is frequently connected to tyrosine kinases (Mahajan and Mahajan, 2015). The accumulation of phosphorylated CHK1, a prime marker of replication fork stalling (Boudny and Trbusek, 2020), occurred in K562 cells that were exposed to HU and MS-275, and such accumulation remained unchanged in the presence of imatinib (**Fig. 5K**).

These data suggest that p73 is a target protein of BCR-ABL in CML cells that are exposed to HU plus MS-275. However, BCR-ABL is detectable in the cytosol whereas p73 is a nuclear protein (Ikeda et al., 2022; Rozenberg et al., 2021). We solved this conundrum by the biochemical fractionation of K562 cells into cytosolic and nuclear fractions. We noted that HU plus MS-275 caused an enrichment of nuclear p73 that was linked to a remarkable nuclear translocation of BCR-ABL (**Fig. 5L**).

We conclude that BCR-ABL is a direct regulator of the phosphorylation of p73 at Tyr99, p73 stabilization, and NOXA induction by p73 upon replication stress and class I HDAC inhibition.

### HDAC1 and HDAC2 safeguard CML cells from DNA replication stress-induced apoptosis via the p73-NOXA axis

Next, we set out to specify which of the HDAC isoforms -HDAC1, HDAC2, or HDAC3- are essential for the survival of CML cells undergoing DNA replication stress. We used the hematopoietic map (HEMAP) database (Pölönen et al., 2019) to evaluate the expression of the mRNAs encoding HDAC1, HDAC2, and HDAC3 in CML cells in relation to normal myeloid cells. Compared to normal myeloid cells (n=690), CML cells (n=234-274) have higher mRNA levels for HDAC1 and HDAC2, but similar levels of HDAC3 (**Figs. 6A** and **S6A**).

To assess if these divergent mRNA expression profiles translate into functional relevance, we applied the 2-thiophenyl biaryl MERCK60 which inhibits HDAC1/HDAC2 selectively (Methot et al., 2008; Stubbs et al., 2015), the fluorinated benzamide RGFP966 which preferentially blocks HDAC3 (Wells et al., 2013), and RNAi against these HDACs to CML cells with DNA replication stress. We found that MERCK60 induced 44% apoptosis in K562 cells and 67% apoptosis in combination with HU after 48 h. These effects and their increase upon the addition of HU were highly significant (**Fig. 5B**). Consistent herewith, MERCK60 induced NOXA (**Fig. 5C**)

We complemented these data with cell cycle analyses by flow cytometry. The overall cell cycle distribution in HU-treated cells showed a highly significant accumulation in S phase which corresponds to DNA replication stress-induced arrest. MERCK60 evokes a moderate G2/M phase arrest at 24 h but does not affect cell cycle stalling by HU. After 24 h, MERCK60 and HU did not significantly raise the apoptotic subG1 population. After 48 h, cells incubated with HU alone or with MERCK60 continued to have significantly less cells in G1 phase than untreated cells. Compared to HU-treated K562 cell populations, K562 cells that were treated with HU and MERCK60 had less cells in S phase. In case of the treatment with HU plus MERCK60, a concurrent rise in the subG1 phase suggests that such cells were lost due to apoptosis. After 48 h, MERCK60 also increased the subG1 phase populations significantly (**Fig. 5D**).

These findings support a model in which HDAC1/HDAC2 inhibition by MERCK60 plus DNA replication stress induction by HU act synergistically to promote apoptosis. We aimed to substantiate these data by silencing HDAC1 or HDAC2 using RNAi. Consistent with the pro-apoptotic impact of MERCK60 (**Fig. 5B**), the combination of RNAi against HDAC1 and HDAC2 induced apoptosis significantly (**Fig. 5E**). K562 control cells that received the scrambled control siRNA did not undergo apoptosis when treated with HU (**Fig. 5E**). In agreement with the synergistic killing effect of HU plus MERCK60 (**Fig. 5B**), RNAi against HDAC1 and HDAC2 induced apoptosis significantly when applied together with HU (**Fig. 5E**). Immunoblotting verified the anticipated depletion of HDAC1/HDAC2 in K562 cells and that both HDACs control the expression of NOXA (**Fig. 5F**). We evaluated if these effects were specific to HDAC1 and HDAC2. In line with the HDAC3 data retrieved from the HEMAP database (Pölönen et al., 2019) in CML cells, we found that inhibition of HDAC3 with RGFP966 or RNAi failed to induce apoptosis or NOXA upon DNA replication stress (**Fig. S6C-E**).

Our findings suggest that HDAC1 and HDAC2 are pharmacological target structures to eliminate CML cells undergoing DNA replication stress.

## Discussion

This study provides multiple lines of evidence for a central pro-apoptotic role of p73 that is induced on the protein level by drugs disturbing DNA replication. P73 binds to the promoter encoding *NOXA* but activates *NOXA* mRNA expression only when HDAC1 and HDAC2 are inhibited pharmacologically or genetically. These findings underscore the importance of p73 as a compensatory mechanism that induces NOXA-dependent apoptosis in p53-deficient CML cells. Consistent with our previous data (Pons et al., 2018; Pons et al., 2021b), we noted that HU stalled CML cells in S phase, without a concomitant induction of apoptosis This makes such cells an ideal model system to analyze HU-induced DNA replication stress, how HDACs regulate this stress response, and if these are targets to break drug resistance. The combined application of the DNA replication stress inducers HU or cytarabine plus the class I HDACi MS-275, FK228, or MERCK60 as well as genetic interference with HDAC1/HDAC2 induces apoptosis in various cultured and primary CML cell types. In contrast, the HDAC6-specific inhibitor marbostat-100 (Sellmer et al., 2018) did not combine favorably with HU against K562 cells, indicating specific protective effects of class I HDACs in cells with DNA replication stress. Using datamining, we found that CML cells overexpress HDAC1 and HDAC2. It is possible that this mechanism evolved in such cells adapt to DNA replication stress that arises during continuous replication of tumor cells and the ensuing insufficient supply of dNTPs.

The BH3-only protein NOXA induces mitochondrial apoptosis (Kale et al., 2018; Morsi et al., 2018) which can explain the highly significant breakdown of mitochondrial membrane integrity. We demonstrate that inhibition of the nuclear HDAC1, HDAC2 sensitizes CML cells undergoing replication stress to NOXA-mediated apoptosis and that p73 acts upstream of NOXA. Therefore, we conclude that these nuclear HDACs act as gatekeepers to prevent DNA replication stress turning into a NOXA-dependent cytotoxic program. Several studies suggest a link between HDACs and NOXA expression but how this can be mechanistically connected target protein BCR-ABL and p73 was unknown (Fischer et al., 2023; Fritsche et al., 2009; Inoue et al., 2008; Liu et al., 2018; Pérez-Perarnau et al., 2011; Sugimoto et al., 2020; Tong et al., 2018; Zhou et al., 2013). Moreover, there is an antagonistic relationship between NOXA and the anti-apoptotic BCL2 proteins MCL1 and BFL-1/A1 (Inoue et al., 2008; Kale et al., 2018; Morsi et al., 2018). These protect mitochondrial membranes to avoid cytochrome c release, caspase-9 and caspase-3 activation, and subsequent apoptotic dismantling of cells (Kale et al., 2018). We show that HU plus MS-275 reduces MCL1. Further anti-apoptotic proteins such as survivin, BCL2, and BCL-XL are also attenuated by MS-275. BAX and BAK, which form pores in mitochondrial membranes for subsequent caspase activation (Kale et al., 2018), remain expressed in HU plus MS-275-treated cells. This also applied to the most pro-apoptotic, short isoforms of BIM. These activate BAX and BAK (Kale et al., 2018). Such datasets illustrate that MS-275 shifts the cellular balance of BCL2 proteins towards pro-apoptotic ones. Macro(autophagy), the self-eating of cells which is usually termed autophagy, has pro-tumorigenic and anti-tumorigenic roles (Shang et al., 2024). While it is known that apoptosis and autophagy interact at various levels (Sarosiek and Wood, 2023; Shang et al., 2024), the molecular mechanisms that connect and integrate these pathways are understood incompletely. Further studies will also show if HDAC1 and HDAC2 control autophagy in CML cells with DNA replication stress. This also applies to a possible accumulation of DNA damage when HDAC1/HDAC2 are inhibited in CML cells with DNA replication stress.

Like p53, p73 contributes to diverse biological processes and cancer hallmarks, such as cell cycle regulation, replicative immortality, and genomic instability. Since the DNA binding domain is conserved within the p53 family, p73 can bind the promoters of the subset of the p53 target genes regulating cell cycle (p21), apoptosis (BAX, FAS, PUMA), and metabolism (Cai et al., 2022; Rozenberg et al., 2021). Under conditions of replicative stress or during S-phase, p73 becomes activated and stabilized through various post-translational modifications (Gong et al., 1999; Yuan et al., 1999), and can then induce the transcription of NOXA. Beyond histones, HDACs also deacetylate a variety of non-histone proteins, including many transcription factors (Mustafa and Krämer, 2023). The acetylation of p73 may also regulate the transcription of the *NOXA* gene. Since the elimination of HDAC1 or HDAC2 increases the levels of NOXA, both HDACs would have p73 as target protein. Such a scenario is reminiscent of the recently found deacetylation of the E3 ubiquitin ligase SIAH2 by both HDAC1 and HDAC2 in myeloproliferative neoplasms (Mustafa et al., 2025).

This work further illustrates that the accumulation of p73 together with increased NOXA levels act as pharmacodynamic markers for pro-apoptotic programs of class I HDACi and DNA replication stress inducers. This highlights the therapeutic potential of targeting p73 pathways to induce cell death in p53-deficient cancers. This conclusion on a predominant role of the p73-NOXA axis in tumors without p53 is supported by the finding that a pharmacological stabilization of p73 by APR-246/PRIMA-1Met triggers NOXA-dependent cell death in cells from esophageal squamous cell carcinoma and multiple myeloma cells lacking functional p53 (Kobayashi et al., 2021; Saha et al., 2013). Previous works disclosed an intricate relationship between tyrosine kinase signaling and DNA damage pathways (Mahajan and Mahajan, 2015; Rozenberg et al., 2021). We reveal that HDAC1 and HDAC2 influence NOXA expression through both chromatin-dependent mechanisms and chromatin-independent mechanisms at the level of tyrosine kinase signaling through BCR-ABL. Furthermore, our findings reveal a critical interaction between BCR-ABL and p73 in CML cells with DNA replication stress and HDAC1/HDAC2 inhibition. We show that BCR-ABL controls the expression and activity of p73, and a subsequent accumulation of NOXA. This aligns with previous findings showing that that nuclear BCR-ABL promotes an accumulation of p73 and the p73-dependent expression of the pro-apoptotic protein PUMA. The latter was found with BCR-ABL that lacks its c-terminal filamentous (F-)actin binding domain (FABD; within the ABL moiety). BCR-ABL^ΔFABD^ accumulates in the nuclear compartment and lacks the ability of BCR-ABL to interact with filamentous actin (Zheng et al., 2024). Studies are underway to address if F-actin controls the cytoplasmic-to-nuclear translocation of BCR-ABL upon DNA replication stress and inhibition of HDACs. Since Tyr99 is not located in the DNA-binding domain of p73 (Cai et al., 2022; Omran et al., 2021), it is unlikely that the phosphorylation of this site affects the chromatin binding ability of p73. This can rather be explained by an HDAC1/HDAC2-dependent chromatin remodeling which modulates the accessibility of specific response elements within the *NOXA* promoter to p73. Binding of p73 activates the expression of *NOXA* mRNA and the accumulation of the pro-apoptotic NOXA protein.

## Conclusion

This work unravels the first data on HDAC1/HDAC2 and BCR-ABL controlling the regulation of the p73-NOXA signaling axis and its functional consequences for CML cells under DNA replication stress. These findings not only enhance our understanding of the cellular response to genotoxic stress in the absence of p53, but also pave the way for novel targeted therapeutic strategies in patients suffering from CML.

## Acknowledgements

We thank Christina Brachetti, Andrea Piée-Staffa, Robert Prause, Birgit Rasenberger, and Anna Frumkina for excellent technical support. This work was mainly funded by the German Research Foundation/Deutsche Forschungsgemeinschaft (KR2291/16-1, project number 496927074). to OHK. Work done in the group of OHK is additionally funded by the projects DFG KR2291/18-1, project number 528202295; KR2291/9-1, project number 427404172; KR2291/12-1, project number 445785155; DFG KR2291/14-1, project number 469954457; KR2291/15-1, project number 495271833; KR2291/17-1, project number 502534123; KR2291/18-1, project number 528202295; funded by the Deutsche Forschungsgemeinschaft (DFG, German Research Foundation) – Project-ID 393547839 – SFB 1361; Project-ID 318346496 - SFB1292 TP21N (MPR); the DAAD Egypt/Germany; the Brigitte und Dr. Konstanze Wegener-Stiftung (project 110); the Walter Schulz-Stiftung; the Hector-Stiftung Weinheim, Germany; and the José Carreras Leukemia Foundation (DJCLS 09 R/2024).

---

***Fig. S1: Inhibition of class I HDACs significantly increases apoptosis in CML cells undergoing replication stress***

A) Flow cytometry was performed to measure annexin-V/PI-positive K562 cells that were treated with 5 µM MS-275 ± 1 mM HU for 24-48 h. Representative dot plots are shown.

B) SubG1 fractions in LAMA-84, MEG-01, KCL-22 and KYO-01 cell cultures that were treated with 5 µM MS-275 ± 1 mM HU for 48 h; n=3, mean±SD; one-way ANOVA, Bonferroni’s multiple comparisons test; p*<0.05, p**<0.01, p***<0.001, p****<0.0001.

C) K562 cells were treated with 1 mM HU ± 5 µM MS-275 for 24-48 h. Exemplary histograms of cell cycle distributions of PI-stained cells are shown.

***Fig S2: Induction of apoptosis during replication stress depends specifically on class I HDAC inhibition***

A) K562 cells were treated with 5 nM FK228 ± 0.5 mM HU for 24 h and 48 h. Flow cytometric analysis was done to count annexin/PI-stained cells; n=3, mean±SD; ANOVA, Bonferroni’s multiple comparisons test; p*<0.05, p**<0.01, p***<0.001, p****<0.0001.

B) Cells treated as indicated in A) were stained with PI for cell cycle distribution analysis; n=3, mean±SD; ANOVA, Bonferroni’s multiple comparisons test; p*<0.05, p**<0.01, p****<0.0001.

C) K562 cells were treated with 0.5 µM marbostat-100 (MARB) ± 1 mM HU for 24 and 48h. Flow cytometric analysis of Annexin/PI-stained cells was performed; n=3, mean±SD; ANOVA, Bonferroni’s multiple comparisons test; ns, not significant.

D) Cell cycle analysis of cells treated as stated in C), n=3, mean±SD; ANOVA, Bonferroni’s multiple comparisons test; p**<0.01, p****<0.0001; ns, not significant.

***Fig. S3: Analysis of additional clones of NOXA^-/-^ and p73^-/-^ K562 cells***

A,B) Wild-type and NOXA knockout (NOXA^-/-^) K562 cells were treated 5 µM MS-275

+0.5 mM HU for 48 h and analyzed for NOXA by immunoblot; HSP90 served as loading control (A) or stained with annexin-V/PI, and subjected to flow cytometry; n=3 ± SD; two-way ANOVA, Bonferroni’s multiple comparisons test; p*<0.05, p**<0.01, p****<0.0001.

C,D) Wild-type and p73 knockout (p73^-/-^) K562 cells were treated 5 µM MS-275 +0.5 mM HU for 48 h and analyzed for NOXA by immunoblot; HSP90 as loading control (A) or stained with annexin-V/PI, and subjected to flow cytometry; n=3 ± SD; two-way ANOVA, Bonferroni’s multiple comparisons test; p*<0.05, p**<0.01, p****<0.0001.

***Fig. S4: p73 and NOXA are induced in K562 cells that are treated with MS-275 plus HU or cytarabine***

A) K562 cells were treated with 1 mM HU and 5 µM MS-275 for 24 h and analyzed for EGR1, MYC, FOXO1, and NFκB p65 expression via immunoblot; β-actin, α-tubulin, and vinculin, loading controls.

B) K562 cells were transfected with the indicated siRNA or plasmids via electroporation.

After 24 h they were treated with 5 μM MS-275 + 1 mM HU for additional 24 h. SubG1 fraction was measured after PI staining via flow cytometry; n=2, mean±SD.

C) K562 cells treated for 24 h with 1 mM HU and 5 µM MS-275 were analyzed for *TP73* mRNA expression via qRT-PCR. Expression was normalized to *GAPDH* and *ACTB* mRNA levels.

D) K562 cells were treated with 5 μM MS-275 ± 1 µM cytarabine. Immunoblot shows p73 after 24 h of incubation and NOXA levels after 48 h of incubation: HSP90, loading. Numbers below immunoblots show densitometric evaluation of relative protein expression. Values were normalized to HSP90; the untreated control is set as 1.

***Fig. S5: HU and MS-275 do not induce NOXA in leukemia cells without BCR-ABL***

A) K562, MV4-11, THP1, U937 cells were treated with 1 mM HU and 5 µM MS-275 for 48h. Cells were lysed and lysates were used to determine protein levels of NOXA. HSP90 serves as loading control.

B) HEL cells were treated with 1 mM HU and 5 µM MS-275 for 24 h and 48 h. Cell lysates were used to determine protein levels of NOXA; β-actin as loading control. Results are representative of at least two independent experiments.

***Fig. S6: Inhibition of HDAC3 does not lead to NOXA expression during replicative stress***

A) The Hemap database (http://hemap.uta.fi/) shows the mRNA expression levels of HDAC3 (n=274) in human CML cell samples compared to normal myeloid cells (n=690); full represents the maximum variation levels.

B) K562 cells were treated with 5 µM RGFP966 ± 1 mM HU for 24-48h. Annexin-V/PI-stained cells were subjected to flow cytometry to measure apoptotic and necrotic cells; n=3, mean±SD; two-way ANOVA, Bonferroni’s multiple comparisons test; no significance.

C) Lysates of cells treated as stated in B) were used to detect the expression of NOXA via immunoblot; a lysate of K562 cells treated with 5 µM MERCK60 for 48 h serves as positive control; HSP90 loading.

D) HDAC3 was knocked down in K562 cells via siRNA. After 24 h, cells were treated with 1 mM HU for additional 24 h. Cell death was measured on the flow cytometer; n=3, mean±SD; two-way ANOVA, Bonferroni’s multiple comparisons test; no significance.

E) Immunoblot shows reduction of HDAC3 by siRNA against HDAC3 and steady NOXA levels in lysates from K562 cells ; HSP90 serves as loading (treatment was done as mentioned in D).

