## Supplementary figures and images for "The Deacetylases HDAC1 and HDAC2 Safeguard BCR-ABL-positive Cells from Replication Stress-Induced Apoptosis via the Nuclear to Mitochondrial p73-NOXA Axis"

### Supplem. Fig. S1-S6

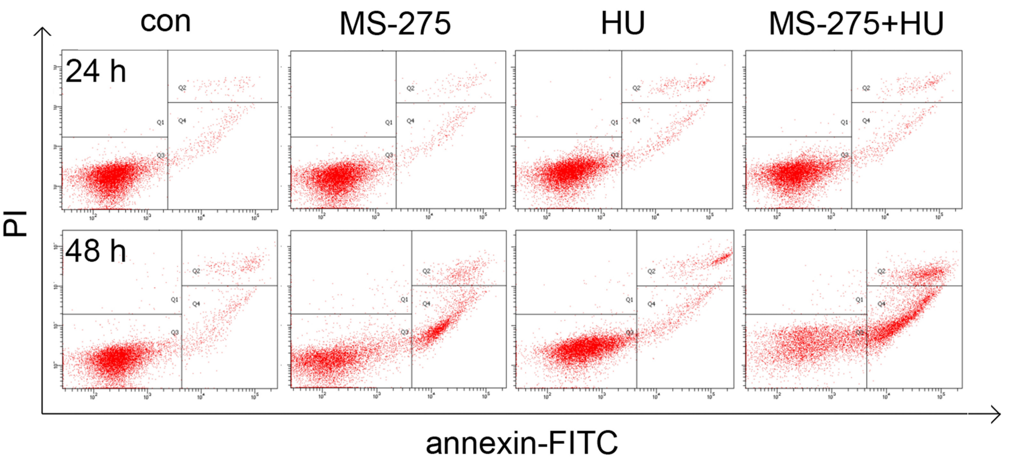

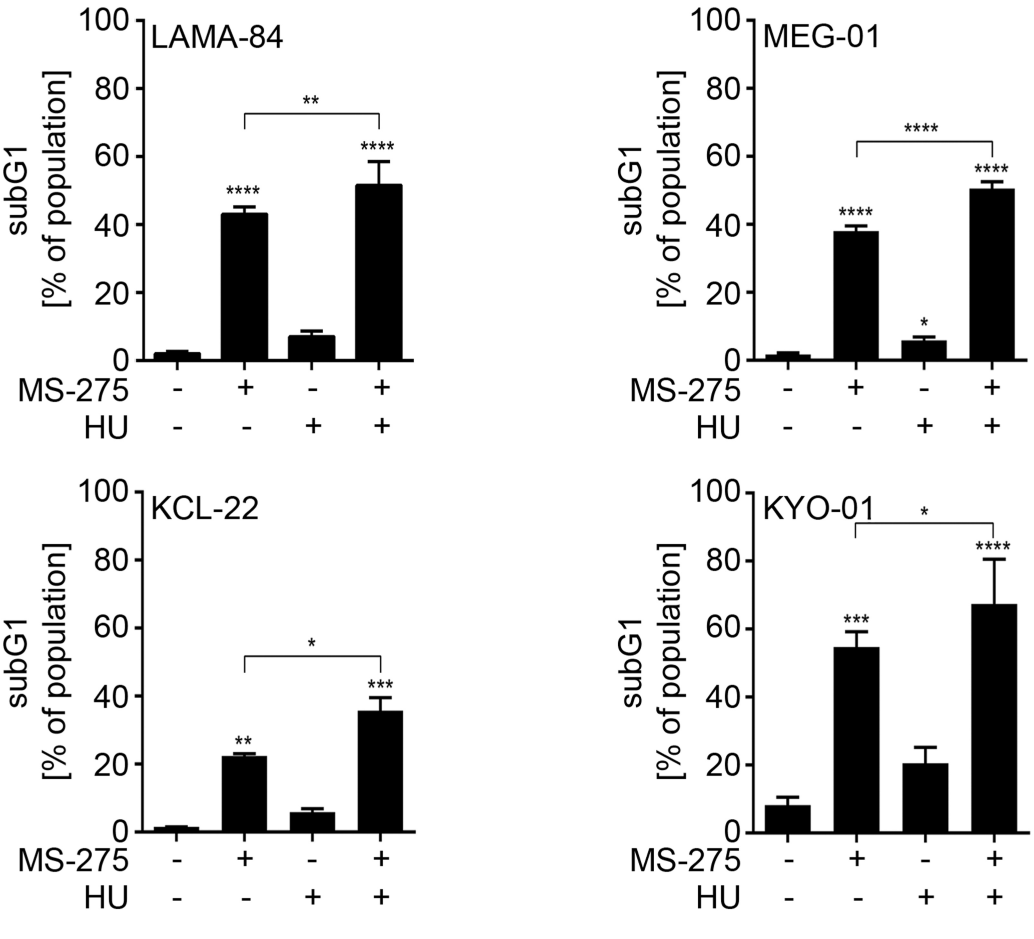

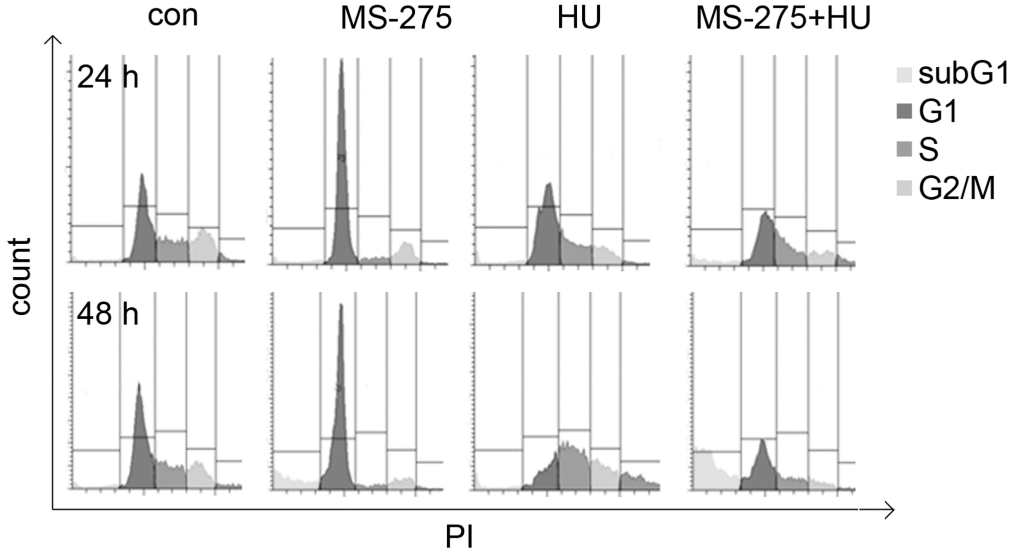

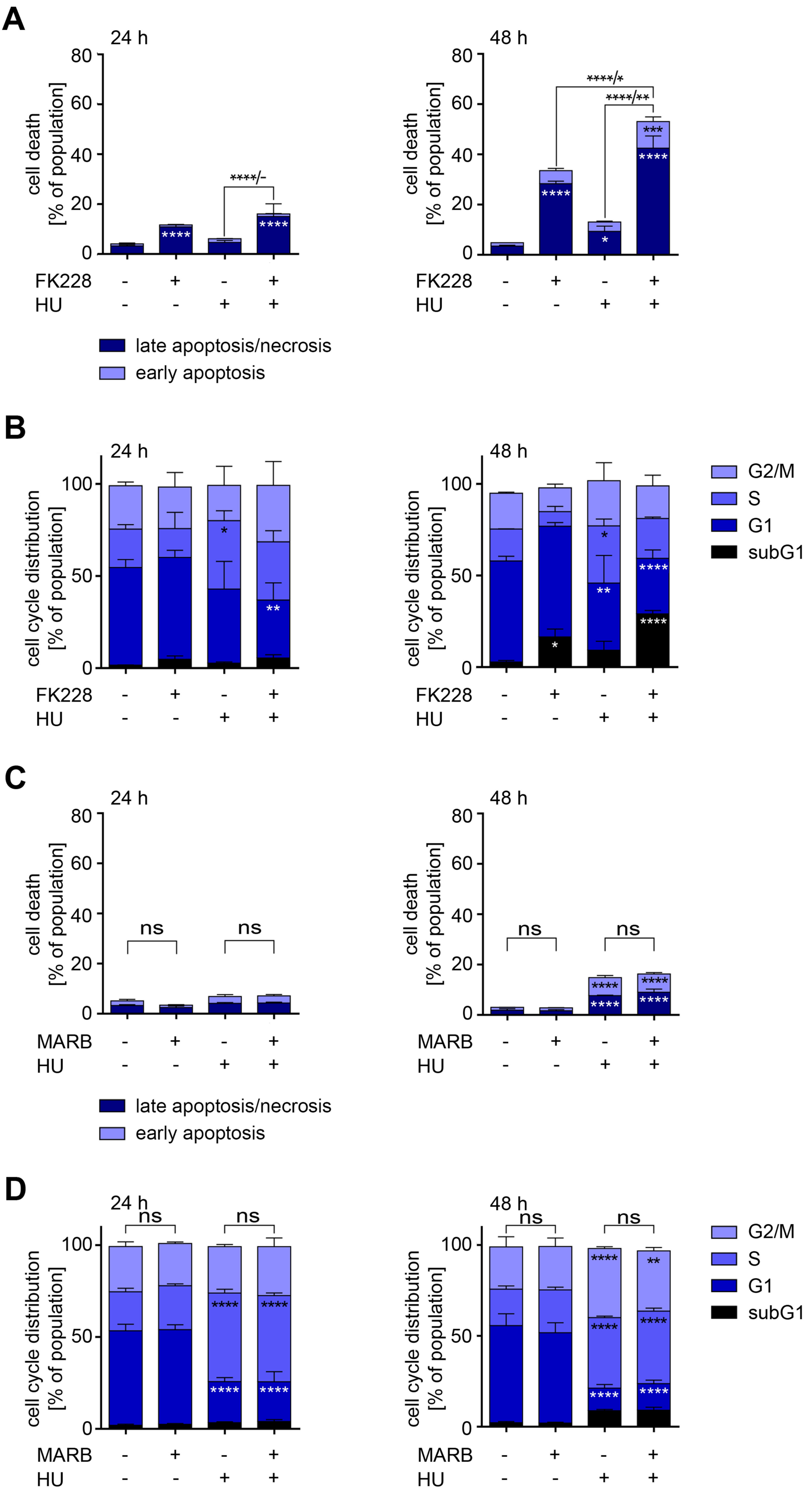

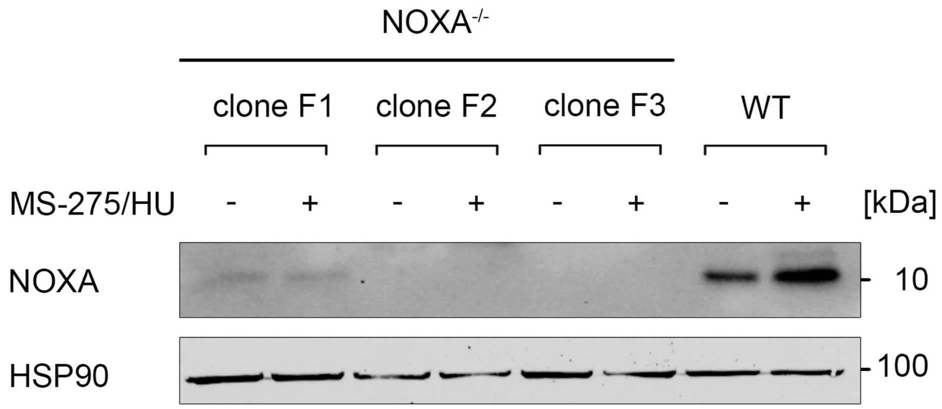

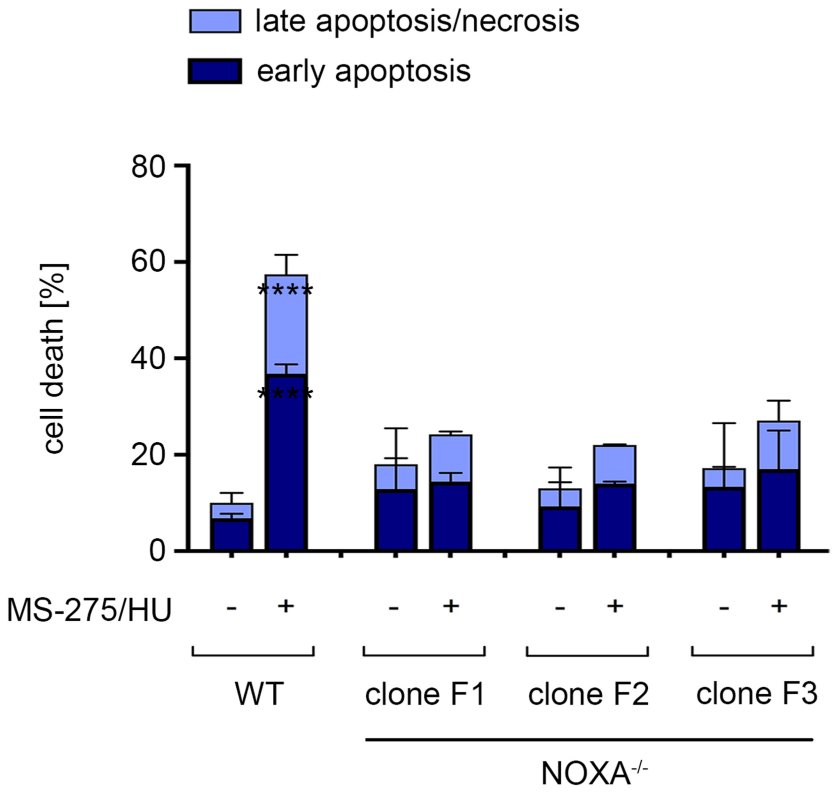

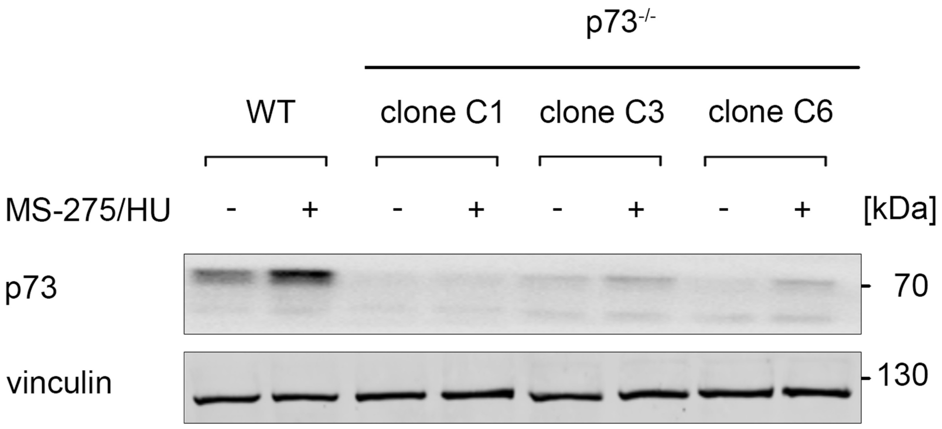

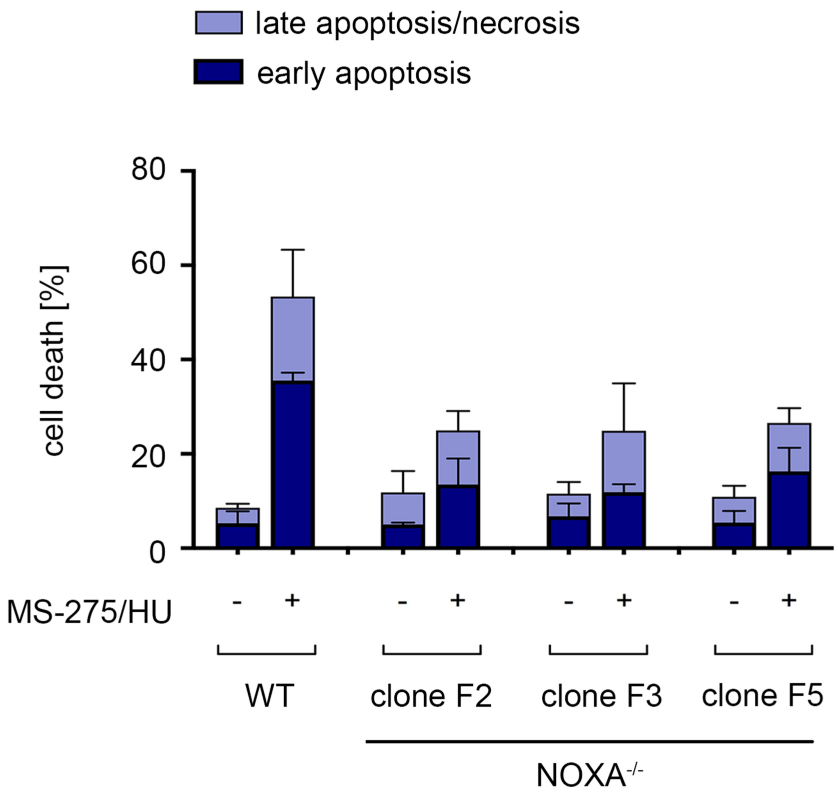

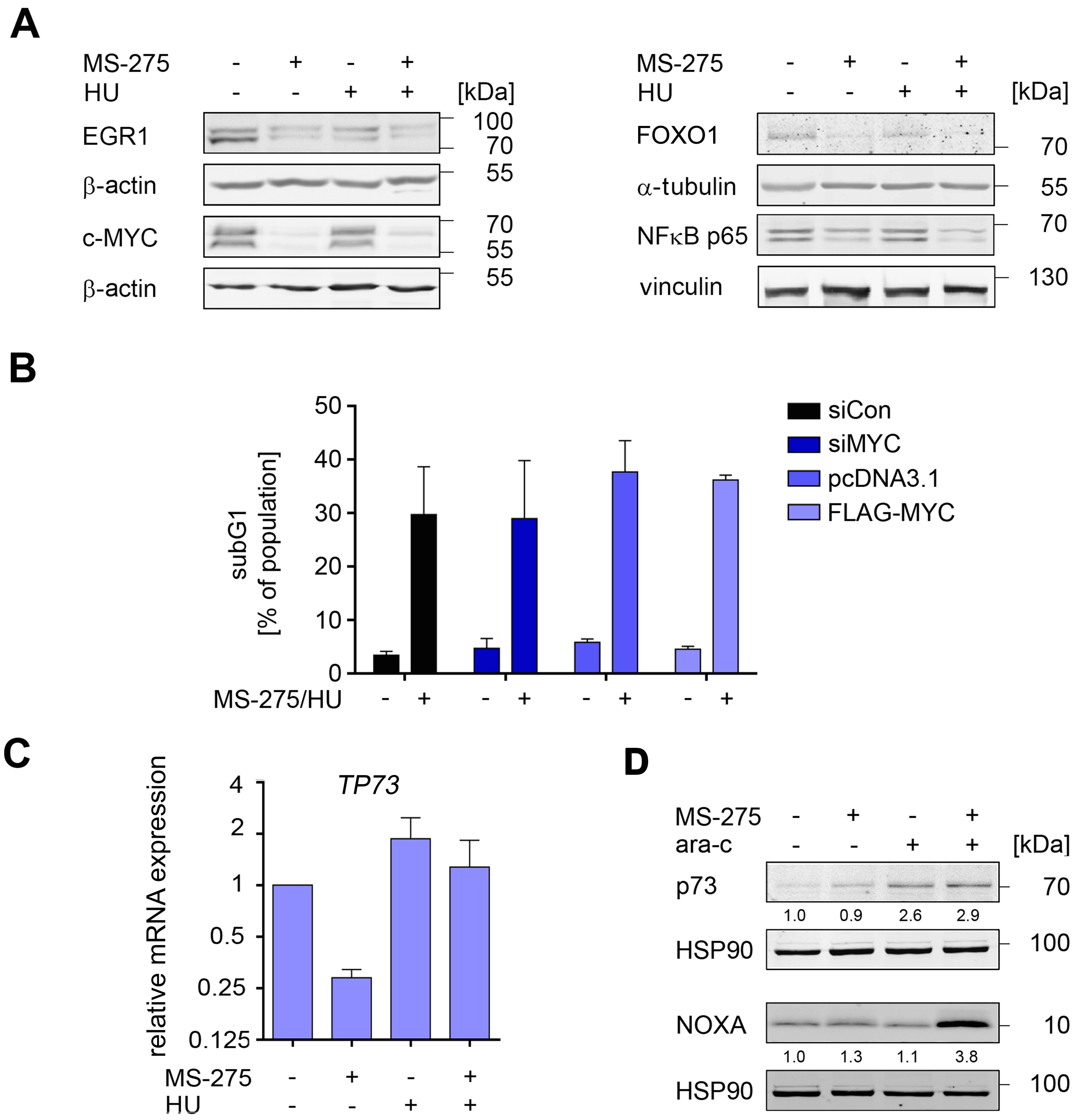

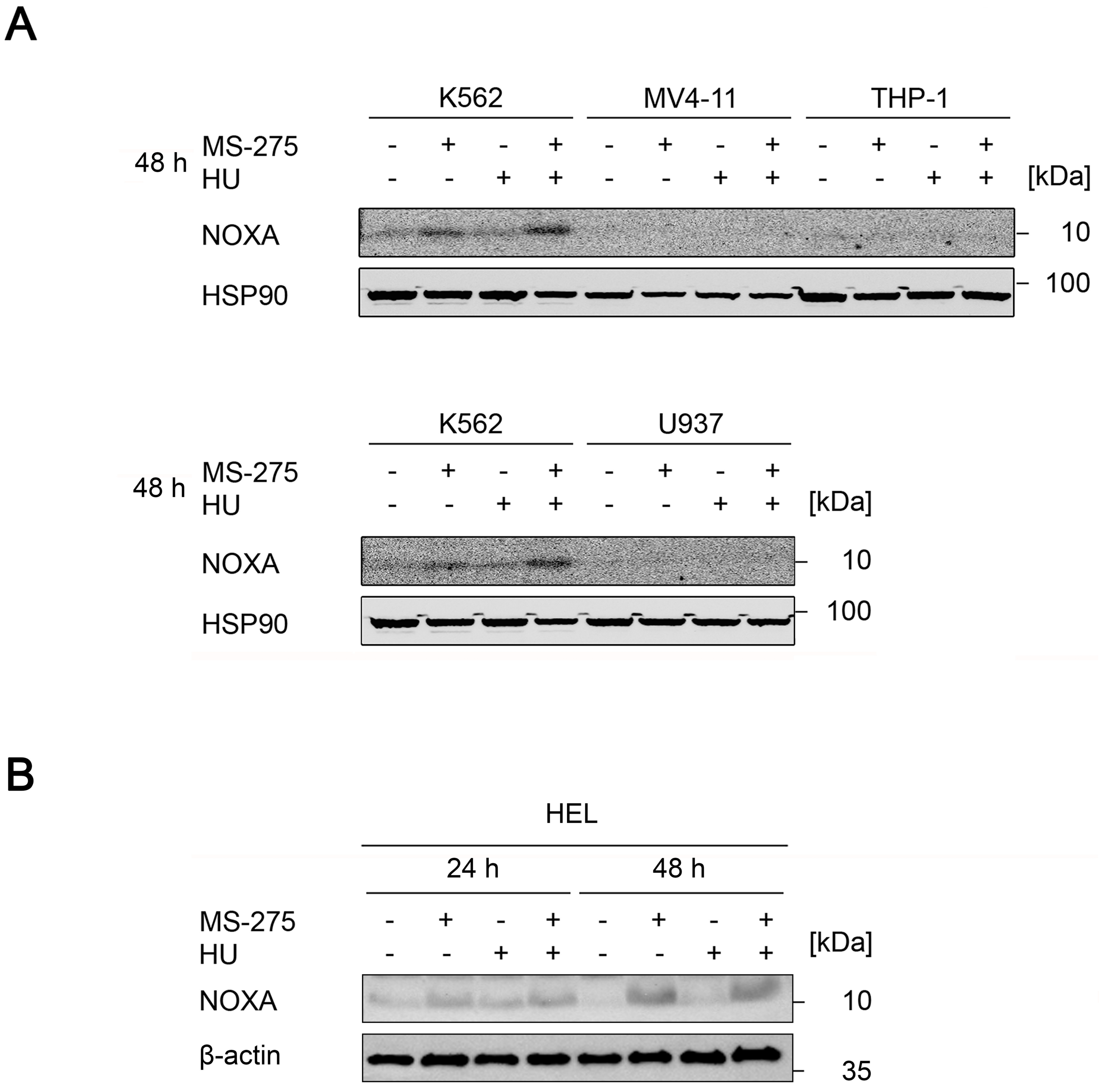

Mustafa et al., Figure S6

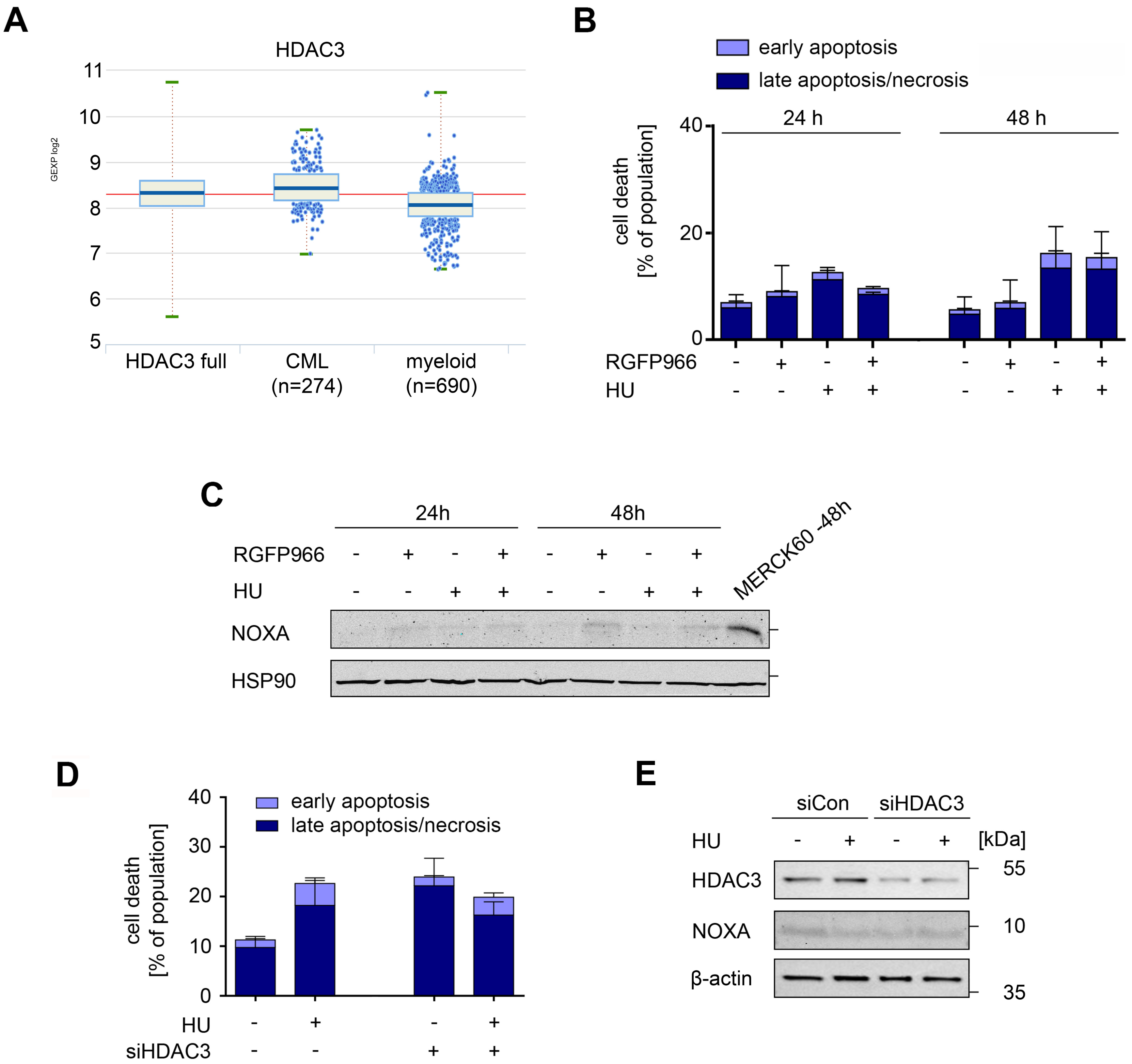
